# Alternative mRNA polyadenylation bridges mitochondrion-to-nucleus communication in Arabidopsis

**DOI:** 10.1101/2022.09.27.509730

**Authors:** Huifang Jia, Weike Zeng, Xiaoyan Zhang, Jiedi Li, Xinyue Bao, Yanming Zhao, Lingjun Zhu, Chongyang Ma, Fengling Wang, Xiangqian Guo, Chun-Peng Song, Liuyin Ma, Xiaohong Zhu

**Affiliations:** State Key Laboratory of Crop Stress Adaptation and Improvement, School of Life Sciences, Henan University, Kaifeng, 475004, China; State Key Laboratory of Cotton Biology, School of Life Sciences, Henan University, Kaifeng, 475004, China; College of Forestry, School of Future Technology, Fujian Agriculture and Forestry University, Fuzhou 350002, China; College of Life Sciences, Fujian Agriculture and Forestry University, Fuzhou 350002, China; Department of Preventive Medicine, Institute of Biomedical Informatics, Academy for Advanced Interdisciplinary Studies, School of Basic Medical Sciences, Henan University, Kaifeng, 475004, China

**Keywords:** Alternative polyadenylation, 3′ UTR shortening, gene expression regulation, translational efficiency, mitochondrial retrograde response, histone demethylation

## Abstract

Mitochondria produce signals besides energy and metabolites that influence plant growth and fitness. However, how mitochondrial signals are relayed to other cellular compartments is largely unknown. By applying poly(A)-site RNA-sequencing (PAS-seq) to wildtype Arabidopsis seedlings and a mutant in the histone demethylase JMJ30 treated with the mitochondrial electron transfer chain inhibitor antimycin A (AA), we identified a previously undefined mitochondrion-to-nucleus communication pathway by which mitochondrial functional state regulates co-transcriptionally alternative polyadenylation (APA) of nuclear mRNA. We observed a global shortening of 3′ untranslated regions (UTRs) as a molecular signature of AA-activated mitochondrial retrograde response (MRR), which contributed in part to translational regulation of auxin response and cell wall biogenesis. JMJ30 regulated AA-induced 3′ UTR shortening, resulting in more transcripts with shortened 3′ UTRs upon AA treatment in a JMJ30 gain-of-function mutant and overexpression lines. We also report on the JMJ30-interacting protein CPSF30, a cleavage and polyadenylation specificity factor that recruits JMJ30 to modulate H3K27me3 status at its target loci. Our study illustrates how epigenetic modification and APA coordinate mitochondrion-to-nucleus communication to allow cells to rapidly respond to changes in mitochondrial functional state and shape plant growth and fitness.

**One-sentence summary:** Epigenetic modification and APA coordinate mitochondrion-to-nucleus communication to allow cells to rapidly respond to changes in mitochondrial functional state and shape plant growth and fitness.

## Introduction

Mitochondria are organelles surrounded by a double membrane, where the electron transport chain (mETC) resides in the inner membrane and generates energy in the form of ATP via oxidative phosphorylation. The activity of mETC-driven oxidative phosphorylation is coupled with the efficiency of the tricarboxylic acid (TCA) cycle taking place in the mitochondrial matrix, thus connecting mETC to cellular metabolism (Meyer et al., 2019; Braun, 2020; Welchen et al., 2021). Mitochondria therefore constitute an important hub for intercompartmental communication. To maintain cellular homeostasis, the signals related to mitochondrial functional status must be conveyed to other cellular compartments to adjust their respective biological processes. Increasing evidence is emerging that perturbation of mitochondrial function is sensed and a response is initiated by the nucleus, the cytosol, chloroplasts, the endoplasmic reticulum (ER), the plasma membrane, and the cell wall, but the underlying communication pathways have not been established (Zhu, 2016; Welchen et al., 2021). The most studied mitochondrial cross-compartment communication is the mitochondrion-to-nucleus retrograde response (MRR), whereby the expression of nuclear genes responds to changes in mitochondrial function (Butow and Avadhani, 2004; Liu and Butow, 2006). Previous studies have identified effectors that mediate this nuclear transcriptional response, using the mETC inhibitor antimycin A (AA) that activates MRR and the nuclear gene *ALTERATIVE OXIDASE1A* (*AOX1A*) as an indicator of the nuclear transcriptional response to AA-induced mitochondrial dysfunction. These effectors are transcription factors: the NAC family members ANAC017 and ANAC013; the WRKY family members WRKY15, WRKY40, and WRKY63; and a subunit of the transcriptional co-activator complex mediator, CDKE1 (Cyclin-dependent Kinase E1) (Vanderauwera et al., 2012; De Clercq et al., 2013; Ng et al., 2013b; Ng et al., 2013a; Van Aken et al., 2013).

In addition to these factors, multiple mutants have been isolated based on MRR defects; their causal mutated genes are related to the establishment of auxin gradients, for example, *BIG*, *PIN-FORMED 1* (*PIN1*), *ATP-BINDING CASSETTE B19* (*ABCB19*), and *ASYMMETRIC LEAVES 1* (*AS1*) (Giraud et al., 2009; Ivanova et al., 2014; Kerchev et al., 2014; Jia et al., 2016; Wang and Auwerx, 2017; Liu et al., 2019; Ohbayashi et al., 2019; Gras et al., 2020). In addition, MRR-defective mutants have been obtained with links to the abscisic acid (ABA) response, such as the causal gene *ABA-INSENSTIVE 4* (*ABI4*) (Giraud et al., 2009). Ethylene signaling is considered an essential regulator of mitochondrial proteostasis (Wang and Auwerx, 2017). Indeed, the exogenous application of ethylene activates the MRR, which is required for ethylene-dependent seed germination (Jurdak et al., 2021). Ethylene and auxin were recently shown to control the MRR via the activation of ANAC017 (He et al., 2022). These studies suggest that plant hormones, in addition to previously proposed candidate signaling molecules like reactive oxygen species (ROS), ATP, and Ca^2+^, convey information from mitochondria to the rest of the cell and/or act as long-distance signals to shape plant growth and development (Berkowitz et al., 2016; Welchen et al., 2021; Welchen and Gonzalez, 2021). However, it is not clear whether or how phytohormone biosynthesis and distribution are linked to changes in mitochondrial function.

Mitochondrion-to-nucleus communication is mostly characterized by transcriptional regulation of nuclear gene expression in response to mitochondrial stress (Butow and Avadhani, 2004; Liu and Butow, 2006). However, it has not been reported in either animals or plants whether and how the co-transcriptional or post-transcriptional regulation of nuclear genes responds to perturbations in mitochondrial function. The co-transcriptional process of alternative polyadenylation (APA) occurs at poly(A) sites (PASs) through the action of the 3′-end processing machinery, resulting in a choice between multiple PASs (Elkon et al., 2013). Thus, APA generates mature transcript variants that differ in the length of their coding sequences and/or their 3′ untranslated regions (3′ UTRs). These mRNA isoforms generated via APA can influence mRNA stability, translation efficiency, mRNA export and localization, as well as protein localization (Tian and Manley, 2017). In fact, APA appears to confer cells the ability to quickly respond to developmental and environmental cues, thus exerting profound effects on cell growth, development, and fitness (Deng and Cao, 2017; Yu et al., 2019b; Zeng et al., 2019; Chakrabarti et al., 2020; You et al., 2021; Ma et al., 2022). Since APA occurs co-transcriptionally, epigenetic factors that regulate nucleosome positioning and chromatin modifications likely affect APA (Zhang et al., 2020; Soles and Shi, 2021). Indeed, in Arabidopsis (*Arabidopsis thaliana*), a study demonstrated that the histone deacetylase HDA6 regulates mRNA polyadenylation, which was supported by a correlation between APA and H3K9ac (acetylation at lysine residue 9 from histone H3) and H3K14ac levels (Lin et al., 2020b). In humans (*Homo sapiens*), the abundance of histone H3K4 and H3K36 methylation is strongly associated with PAS and likely affects the choice of PAS, although their regulatory role in APA remains unknown (Khaladkar et al., 2011; Lee and Chen, 2013). In budding yeast (*Saccharomyces cerevisiae*), the histone H3K4 methyltransferase Set1 and the histone H3K36 methyltransferase Set2 were shown to control PAS choice, likely via repressing the phosphorylation of the C-terminal domain of RNA polymerase II (RNAPII-CTD), thus blocking the recruitment of the 3′-end processing complex. Moreover, both Set1 and Set2 regulate APA induced by nutritional stress, indicating that APA regulation mediated by histone demethylases responds to stress stimuli (Michaels et al., 2020).

Methylation of histone H3 at lysine residues is an important chromatin modification mark for the epigenetic regulation of transcription (Liu et al., 2010; Cheng et al., 2020). Tri-methylation of histone H3 at lysine 27 (H3K27me3) is often associated with low-expressed genes and is thus considered a transcriptional repression mark (Zhang et al., 2007; Xiao et al., 2016). In plants, the chromatin of many developmental genes and stress-responsive genes is decorated by H3K27me3, suggesting a potential epigenetic regulation of plant development and stress adaptation through histone H3K27me3 modifiers (Lu et al., 2011; Li et al., 2013; Gan et al., 2014; Zheng et al., 2019; Cheng et al., 2020; Wang et al., 2021). The H3K27me3 mark is dynamically deposited by SET (Su(Var), E(Z), Trithorax) domain–containing histone methyltransferases and removed by Jumonji C (JmjC) domain–containing histone demethylases. In Arabidopsis, five JMJ demethylases exhibit H3K27me3 demethylation activity (Lu et al., 2008). JMJ30 (encoded by At3g20810) is a member of the JMJD6 group of JMJ proteins and controls H3K27me3 levels over the flowering repressor locus *FLOWERING LOCUS C* (*FLC*) and over the *HEAT SHOCK PROTEIN* (*HSP*) locus to maintain heat stress memory (Gan et al., 2014; Yamaguchi et al., 2021). These results indicate that JMJ30 functions in reprograming the H3K27me3 status at specific loci during plant development and in response to environmental signals.

Here, we tested the hypothesis that APA and epigenetic factors mediate mitochondrion-to-nucleus communication. To this end, we examined APA events in response to AA-induced mitochondrial stress in wildtype Arabidopsis and *jmj30* mutant seedlings. We determined that AA induces a global shortening of 3′ UTRs and that *JMJ30* expression enhances the number of genes whose 3′ UTRs shorten in response to AA treatment. We also identified the cleavage and polyadenylation specificity factor CPSF30 as an interactor of JMJ30. Using a chromatin immunoprecipitation (ChIP) assay, we demonstrate that JMJ30 controls the H3K27me3 status at CPSF30 target loci, supporting a model whereby CPSF30 facilitates the recruitment of JMJ30 and the co-action of CPSF30 and JMJ30 not only regulates 3′-end processing via potential effects on PAS usage and/or efficiency of the 3′-end processing machinery, but also controls the transcription of nuclear genes in response to mitochondrial functional state.

## Results

### Mitochondrial stress elicits global changes in alternative polyadenylation

To explore how nuclear transcriptional and co-transcriptional processes respond to the functional status of mitochondria, we performed transcriptome deep sequencing (RNA-seq) and poly(A)-test RNA-sequencing (PAT-seq) using total RNA extracted from 10-d-old Arabidopsis wildtype (Col-0) seedlings treated with antimycin A (AA) or treated with an equivalent volume of DMSO (AA solvent, as control) (Supplemental Figure S1 A and B). AA treatment elicits mitochondrial stress and activates the MRR by inhibiting the activity of the mitochondrial electron transport chain (mETC) (Wikstrom and Berden, 1972; Slater, 1973; Huang et al., 2005). It was reported that AA also inhibits cyclic electron transport in plastids (Antal et al., 2013; Taira et al., 2013; Takagi et al., 2019). To check if AA treatment affects the plastid function, we examine plastid retrograde marker gene expression under the conditions applied to trigger MRR. The result showed that the expression of plastid retrograde marker genes is not significantly altered by AA treatment (Supplemental Figure S1C) while lincomycin (Lin) and norflurazon (NF) that are known to induce chloroplast retrograde response significantly inhibit the marker gene expression (Supplemental Figure S1D). We conclude that AA has no major impacts on plastid function under the conditions used in this study.

To determine the effects of AA-induced mitochondrial stress on mRNA 3′-end processing, we examined the global distribution of poly(A) sites (PASs). We identified 68,588,124 PASs reads and 55,427,354 of them were uniquely mapped to Arabidopsis genome (Supplemental Data Set S1: T1). After removing 1,076,651 internal priming reads (PASs reads mapped to genomic regions with six or more consecutive Adenines at their terminal), 48,564,521 qualified PASs reads were obtained (Supplemental Data Set S1:). Adjacent poly(A) sites within 24 nt were grouped into poly(A) site clusters (PACs) to reduce the effects of microheterogeneity and a total of 59,872 PACs were identified. As expected, PACs were predominantly (∼90%) mapped to 3′ UTRs of coding genes, with a much smaller fraction mapping to 5′ UTRs, exons, introns, and intergenic regions. About 60% of genes had at least two poly(A) sites (Supplemental Figure S1E). We observed no significant difference in PAS distribution between control and AA-treated seedlings (Figure 1A). However, we did notice a relative shift in PAS position in response to AA treatment. We detected 2,613 genes that displayed a switch in their PAS from a distal (dPA) to a proximal (pPA) site (Figure 1B, red dots). Conversely, we identified 957 genes (blue dots) showing a shift in their PAS from a proximal (pPA) to a distal (dPA) site. Notably, this number was significantly smaller than genes exhibiting a distal-to-proximal shift (Chi-Square Test, *P* = 3.19×10^−129^). We visually inspected mapped PAS-seq reads in the Integrated Genome Viewer (IGV) browser for two selected genes, *INDOLE-3-ACETIC ACID INDUCIBLE 9* (*IAA9*) and *PROTEIN PHOSPHATASE 2C* (*PP2C*), in auxin and ABA signaling, respectively, which demonstrated that AA treatment results in both a shift and a change in the number of reads in ploy(A) sites (Figure 1C). Together, these results clearly indicated that AA treatment affects 3′ UTR processing and elicits global alternative polyadenylation (APA).

**Figure 1.**
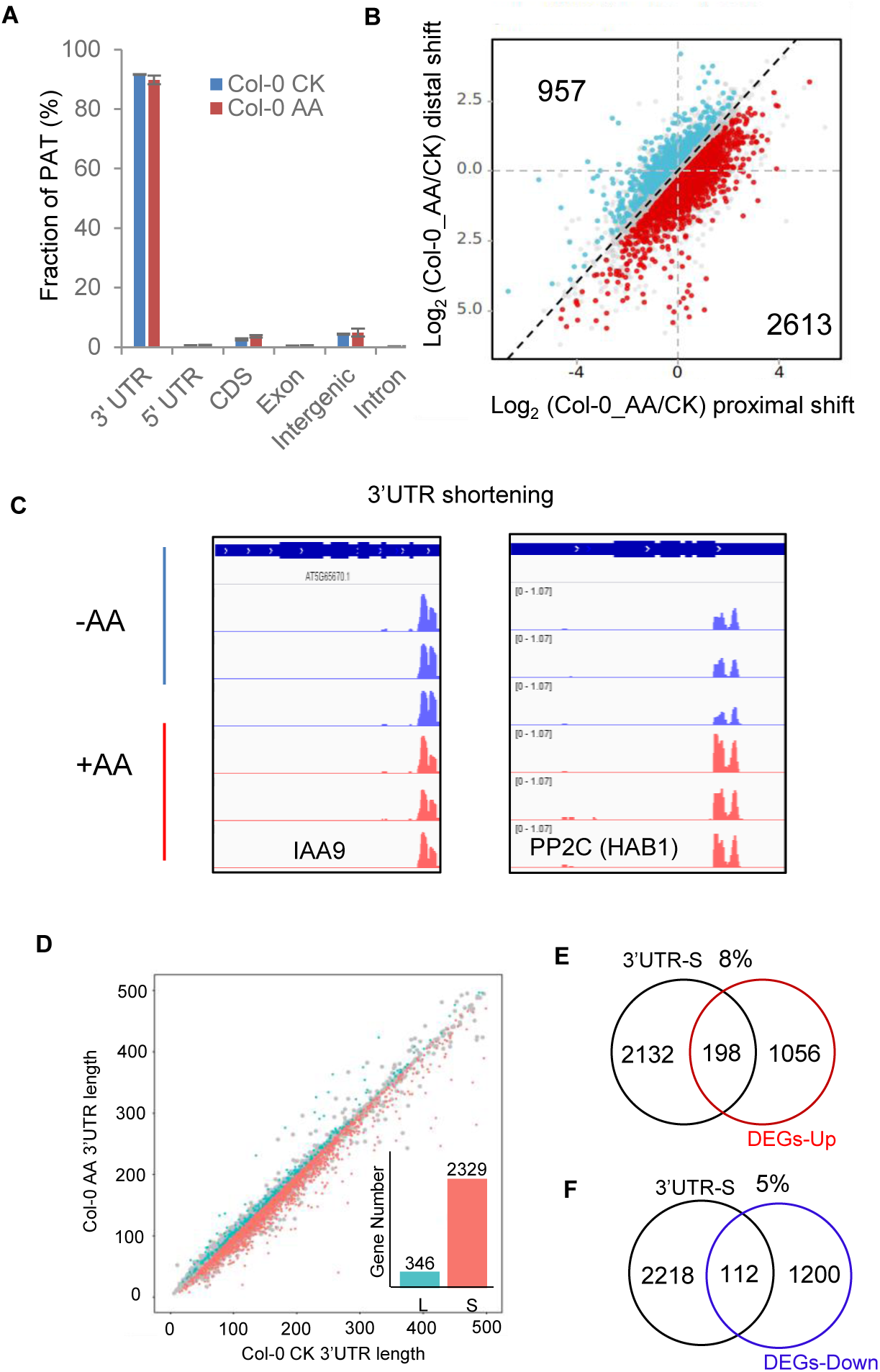
Antimycin A–induced mitochondrial stress leads to global alternative polyadenylation. **A**, Antimycin A (AA) treatment has no significant effect on the distribution of poly(A) tails (PATs) across genomic regions. **B**, AA induces a global shift in poly(A) sites. The scatterplot shows distal-to-proximal shifts (2,613 genes, red dots) and proximal-to-distal shifts (957 genes, blue dots) in poly(A) sites. **C**, IGV browser window showing AA-induced 3′ UTR shortening. *IAA9* and *PP2C* from the auxin and ABA pathways, respectively, show 3′ UTR shortening in response to AA treatment. **D**, The shortening of 3′ UTRs predominates in AA-treated samples. Scatterplot of weighted 3′ UTR lengths for transcripts from AA-treated and control Col-0 seedlings. L, 3′ UTR lengthening; S, 3′ UTR shortening. The inset shows the number of transcripts with L and S. **E** and **F**. Venn diagrams showing the extent of overlap between AA-induced 3′ UTR shortening (UTR-S) and upregulated genes (**E**) or downregulated genes (**F**). See also Supplemental Data Set S1.

Over 80% of all PASs are located in the 3′ UTR regions of mRNA (Wu et al., 2011b). We therefore determined the extent of variation in 3′ UTR length between AA-treated and control seedlings. We divided genes in terms of their 3′ UTR length based on the criteria r < 0 for 3′ UTR shortening and r > 0 for 3′ UTR lengthening (*P* < 0.05) and observed that 3′ UTR shortening events (2,329) are far more abundant than 3′ UTR lengthening events (346, Chi-Square Test, *P* = 6.30×10^−259^) in AA-treated wildtype seedlings (Figure 1D, Supplemental Data Set S1: T2). These results indicated that most transcripts use the proximal PAS to terminate early during polyadenylation in response to AA treatment. The AA-induced transcriptome was thus characterized by the predominance of mRNA isoforms with shortened 3′ UTRs. We also investigated whether 3′ UTR shortening upon AA treatment correlates with differential gene expression. A total of 71,557,944 RNA-seq reads were uniquely mapped to Arabidopsis genome (Supplemental Data Set S1: T3) and 2330 genes were differential expressed genes upon AA treatment compared to wildtype control (Supplemental Data Set S1: T4-5). Notably, we detected no strong correlation between 3′ UTR shortening and the relative transcript levels of DEGs. Although 8% (198/2,330) and 5% (112/2,330) of the transcripts whose 3′ UTR was shortened exhibited a correlation with higher or lower expression levels, respectively (310, Chi-Square Test, *P* = 2.24×10^−6^ Figure 1, E and F), most (87%) transcripts showed no change in their expression upon AA treatment (Supplemental Data Set S1:T6-T7). Together, these results strongly suggest that the 3′ UTR processing of transcripts derived from nuclear genes responds to mitochondrial stress and that AA imposes a preferred usage for the proximal PAS in the 3′-most coding exon of mRNAs, resulting in the shortening of 3′ UTRs.

### 3′ UTR shortening contributes to translational regulation of nuclear genes in phytohormone and cell wall biogenesis pathways upon AA-induced mitochondrial stress

The lack of strong correlation between 3′ UTR shortening and relative transcript levels prompted us to examine whether 3′ UTR shortening might affect mRNA translation. We first analyzed Ribo-seq data, performed for AA-treated and control samples. AA treatment caused a clear shift of mRNAs from polysomes to monosomes (Figure 2A). This result indicated that AA-induced mitochondrial stress causes a global repression of cytosolic translation. To measure translation efficiency (TE), we generated RNA-seq libraries of the monosomal and polysomal RNA fractions from AA-treated and control samples. We applied a cutoff for TE of |Log_2_(FC)| > 0.32 (FC: AA_polysome/CK_polysome vs AA_monosome/CK_monosome) and a variable influence on projection (VIP) value (of either monosome or polysome comparisons) > 1, yielding 4,970 and 4,205 transcripts with significantly higher and lower TE, respectively (Supplemental Data Set S2: T1-T2). Notably, transcripts with shortened 3′ UTRs contributed significantly to the repression of cytosolic translation (Figure 2B, Supplemental Data Set S2: T3-4). About ∼16% (356/2,330) and 20% (458/2,330) of 3′ UTR–shortened transcripts are associated with high or low TE of transcripts (Figure 2, C and D), which is much higher than the percentage associated with DEGs (8% and 5%, Figure 1, E and F). These results suggest that 3′ UTR–shortened transcripts contribute more to post-transcriptional regulation.

**Figure 2.**
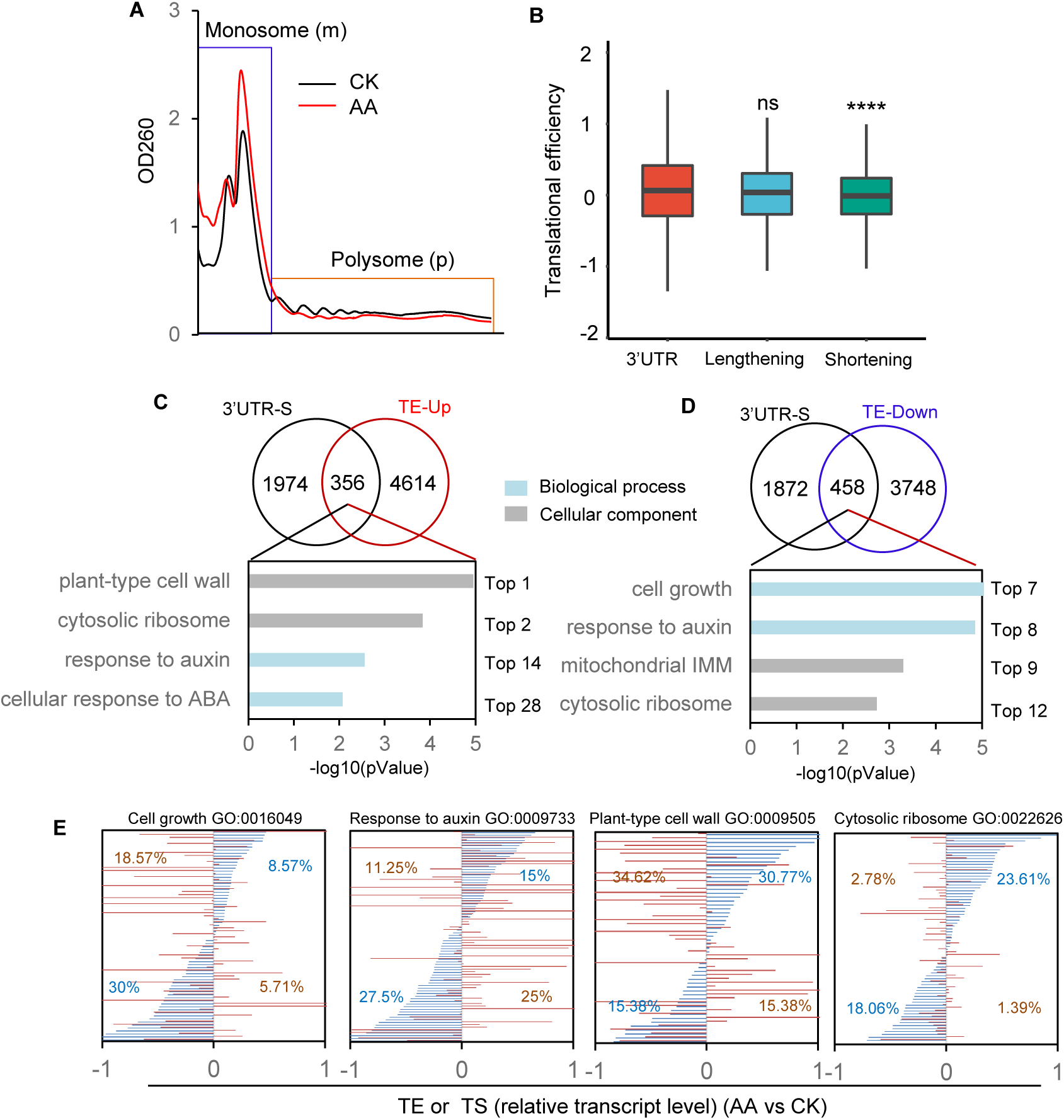
3′ UTR shortening contributes to cytosolic translational regulation of nuclear genes in response to mitochondrial stress. **A**, Representative polysome profiles showing the alteration of cytosolic polysome abundance in AA-treated samples (red line) compared to untreated samples (black line). RNAs isolated from highly translated fractions (polysomes, shown in the red box) and lesser translated fractions (monosomes, shown in the blue box) were used for RNA-seq analysis. **B**, 3′ UTR shortening significantly contributes to a decrease in translational efficiency. **C**, GO enrichment analysis of genes whose transcripts undergo 3′ UTR shortening upon AA treatment and exhibit altered translation efficiency. The Venn diagrams show the extent of overlap between genes displaying AA-induced 3′ UTR shortening and AA-induced upregulated translation efficiency (TE) (**C**) or downregulated TE (**D**). The enriched biological processes are indicated below each diagram. See also a complete list of enriched terms in Supplemental Data Set S2. E, Visualization of TE (blue lines) and transcript abundance (red lines) of genes whose transcripts exhibit 3′ UTR shortening enriched in four biological processes. GO terms are cell growth (GO:0016049), response to auxin (GO:0009733), plant-type cell wall (GO:0009505), and cytosolic ribosome (GO:0022626). The numbers within each panel are the percentages of significant hits that are defined by the criteria of |Log_2_(FC)| ≥ 1 and *P*-adj < 0.05 for transcripts and |Log_2_FC| > 0.32, VIP (of either mono or poly comparisons) > 1 for TE. See also a complete list of genes in Supplemental Data Set S2.

To explore whether the 3′ UTR shortening–mediated translational regulation is a mitochondrial stress adaptive mechanism, we performed a Gene Ontology (GO) analysis of 3′ UTR–shortened transcripts that have either high TE (Figure 2D, Supplemental Data Set S2: T5) or low TE (Figure 2E, Supplemental Data Set S2: T6). We observed strong enrichment for the GO terms “plant-type cell wall” (GO:0009505) and “cellular response to ABA” (GO:0071215) among 3′ UTR–shortened transcripts with high TE, and “cell growth” (GO:0016049) and “mitochondrial inner membrane” (GO:0005743) for 3′ UTR–shortened transcripts with low TE. In addition, the GO terms “cytosolic ribosome” (GO:0022626) and “response to auxin” (GO:0009733) were both enriched in transcripts with high TE or low TE, although with different enrichment rankings (Figure 2, C and D; Supplemental Data Set S2:T5-T6). We plotted TE and transcript levels (TS) of 3′ UTR–shortened transcripts enriched in the four biological processes and cellular components listed above (Supplemental Data Set S2: T7). As shown in Figure 2E, of the 3′ UTR–shortened transcripts enriched in the categories “cell growth” and “response to auxin”, 30% and 27.5% (significant hits), respectively, exhibited low TE. Interestingly, most 3′ UTR–shortened transcripts associated with the cell wall were transcriptionally downregulated (34.6% significant hits), but showed high TE (30.8% significant hits). The levels of 3′ UTR–shortened transcripts related to cytosolic ribosomes displayed no significant change, but half showed either high TE (23.6% significant hits) or low TE (18.1% significant hits) (Figure 2E). These results suggest that 3′ UTR shortening counteracts or synergizes transcriptional and translational regulation and potentially determines protein production. For 3′ UTR–shortened transcripts with no significant changes in their transcript levels, for example, cytosolic ribosomes, translational efficiency likely determines protein production. Thus, 3′ UTR shortening mediates transcriptional and translational regulation of nuclear genes that are involved in cytosolic translation, phytohormone responses, cell wall biogenesis and cell growth.

We further manually examined selected transcripts from each GO category. For cell growth– related genes, we selected the cell wall–related genes *ACTIN2* (*ACT2*), *ACT7*, *KDSA1* (encoding 3-DEOXY-D-MANNO-OCTULOSONATE 8-PHOSPHATE SYNTHASE), *EXPA8* (*EXPANSIN A8*), *XTH4* (*XYLOGLUCAN ENDOTRANSGLUCOSYLASE/HYDROLASE 4*), *MUR1* (*MURUS1*, encoding a GDP-D-MANNOSE-4,6-DEHYDRATASE 2), and *ARL* (*ARGOS-LIKE*, encoding a downstream regulator of BRASSINOSTEROID-INSENSITIVE 1 [BRI1]), which all showed low TE (Supplemental Figure S2A, Supplemental Data Set S3). Auxin-responsive genes, like the auxin receptor *TIR1* (*TRANSPORT INHIBITOR RESPONSE 1*), *SAUR1* (*SMALL AUXIN UPREGULATED RNA*), *SAUR30*, *SAUR77*, *IAA9*, *IAA19*, and *GH3.17* (*GRETCHEN HAGEN 3.17*, encoding an IAA-amido synthase) all had low TE (Supplemental Figure S3A). Transcripts for mitochondrial inner membrane components including several subunits of mitochondrial ATP synthase, AAC1 (ADP/ATP CARRIER 1), DIC2 (DICARBOXYLATE CARRIER 2), and SAMC2 (S-ADENOSYLMETHIONINE CARRIER 2), or mitochondrial import (TIM50, TRANSLOCON OF THE INNER ENVELOPE OF MITOCHONDRIA) exhibited low TE (Supplemental Figure 2SB). Most proteins that generate precursor metabolites and energy are plastid-localized proteins, including PSAN (only subunit of photosystem I [PSI]), ATPD (chloroplast ATPase delta-subunit), FNR1 (FERREDOXIN-NADP(+)-OXIDOREDUCTASE 1), GAPB (GLYCERALDEHYDE-3-PHOSPHATE DEHYDROGENASE), PGK1 (PHOSPHOGLYCERATE KINASE 1), PSBW (a protein similar to the PSII reaction center subunit W), PSBP-1 (a 23-kD extrinsic protein that is part of PSII), and TLP18.3 (THYLAKOID LUMEN PROTEIN 18.3 regulating the PSII repair cycle). Their transcripts all had shortened 3′ UTRs and low TE (Supplemental Figure S2B). Cytosolic ribosome transcripts with shortened 3′ UTRs showed little change in their transcript level but were equally divided into high-TE or low-TE transcripts (Supplemental Figure S2C). Key genes involved in cell wall biogenesis, for example, *CESA2* (*CELLULOSE SYNTHASE A2*) and *CESA5* in cellulose biosynthesis; *PAE11* (*PECTIN ACETYLESTERASE 11*), *PME5* (*PECTIN METHYLESTERASE 5*), and *PME46* in pectin remodeling; and *EXT8* (*EXTENSIN 8*), *EXT12*, and *EXT13* in cell extension, showed AA-induced high TE but low transcript levels (Supplemental Figure S2D). The key genes encoding regulators of the ABA signaling pathway, such as ABF3 (ABA-RESPONSIVE ELEMENTS-BINDING FACTOR 3), HAB1 (HYPERSENSITIVE TO ABA1), ABO3 (ABA-OVERLY SENSITIVE 1), and ARCK1 (ABA- AND OSMOTIC-STRESS-INDUCIBLE RECEPTOR-LIKE CYTOSOLIC KINASE1), displayed high TE and high TS (Supplemental Figure S2D). Together, these results suggest that 3′ UTR shortening contributes to cytosolic translational regulation of plant hormone response, cell growth, cell wall biogenesis, and mitochondrion- and chloroplast-mediated energy production in response to mitochondrial stress.

### The histone demethylase JMJ30 regulates 3′ UTR processing in response to mitochondrial stress

To identify effectors controlling 3′ UTR processing in response to mitochondrial stress, we analyzed the PAS-seq data obtained from AA-treated and control seedlings for the wildtype Col-0, a histone demethylase JMJ30 mutant (*jmj30-1*, SAIL_811_H12), and a *JMJ30* overexpression line (*JMJ30OX*) (Supplemental Figure S1, A and B). Surprisingly, relative *JMJ30* transcript levels were about 8-fold higher in *jmj30-1* and 4,000-fold higher than those of Col-0, indicating that *jmj30-1* is a gain-of-function mutant (Supplemental Figure S3). Similar to Col-0, we detected many transcripts undergoing PAS switching from dPA to pPA in the *jmj30-1* mutant (Supplemental Figure S4A; 2,665 transcripts; Supplemental Data Set S3:T1-2) and *JMJ30OX* (Supplemental Figure S4B; 2,969 transcripts Supplemental Data Set S3:T3) upon AA treatment, with fewer transcripts showing PAS switching from pPA to dPA (722 transcripts in the *jmj30-1* mutant and 496 transcripts in *JMJ30OX* seedlings; Supplemental Figure S4, A and B). We applied the same criteria as above in Col-0 seedlings, which indicated that 3′ UTR shortening events are much more frequent than lengthening events. In addition, we detected fewer lengthening events in the *jmj30-1* mutant and *JMJ30OX* seedlings than in Col-0 (Figure 3A), suggesting that JMJ30 controls 3′ UTR shortening in response to mitochondrial stress. A cross-analysis of transcript levels and 3′ UTR index suggested that 3′ UTR shortening, but not 3′ UTR lengthening, significantly contributes to the upregulation of gene expression in *jmj30-1* (Figure 3B) and *JMJ30OX* (Figure 3C). We also investigated to what extent 3′ UTR shortening correlated with differential gene expression upon AA treatment of *jmj30-1* and *JMJ30OX* seedlings by extracting DEGs in response to AA treatment and plotting the fold-change in transcript levels as a function of 3′ UTR length index. We identified 2,297 (*jmj30-1*) and 3,267 (*JMJ30OX*) DEGs (Supplemental Data Set S3:T4-8). We observed no strong correlation between 3′ UTR length and fold-change in expression levels for either *jmj30-1* or *JMJ30OX*. Among the transcripts with shortened 3′ UTRs, we detected relatively more upregulated DEGs (11% in *jmj30-1* and 14% in *JMJ30OX*) and fewer downregulated DEGs (4% in *jmj30-1* and 3% in *JMJ30OX*) (Supplemental Figure S4, C and D). These results indicated that transcripts with shortened 3′ UTRs upon AA treatment contribute more to the upregulation of DEGs as *JMJ30* levels increase. We also identified 819 transcripts with shortened 3′ UTRs following AA treatment common to Col-0, *jmj30-1*, and *JMJ30OX*, as well as over 800 unique transcripts to each genotype, suggesting that a group of 3′ UTR shortening events can be specifically attributed to JMJ30 function (Supplemental Figure S4H). We also observed this uniqueness in the different outcomes of GO term enrichment analysis. The top enriched biological processes associated with 3′ UTR–shortened transcripts in Col-0 (Supplemental Figure S4E) were distinct from those obtained with similar transcripts in *jmj30-1* (Supplemental Figure S4F) and *JMJ30OX* (Supplemental Figure S4G). We focused on the top enriched processes “response to auxin”, “response to ABA”, “plant-type cell wall”, and “cytosolic ribosome” to determine the different processes that JMJ30 function may contribute to. We detected more genes enriched in “cytosolic ribosome”, “plant-type cell wall”, and “response to ABA” but much fewer genes enriched in “response to auxin” in *JMJ30OX* compared to Col-0, prompting us to hypothesize that the degree of enrichment in these biological processes is related to *JMJ30* expression levels (Figure 3D). Although some 3′ UTR–shortened transcripts were common to the three genotypes, a fraction was unique to *jmj30* and *JMJ30OX* in four biological processes (Figure 3D).

**Figure 3.**
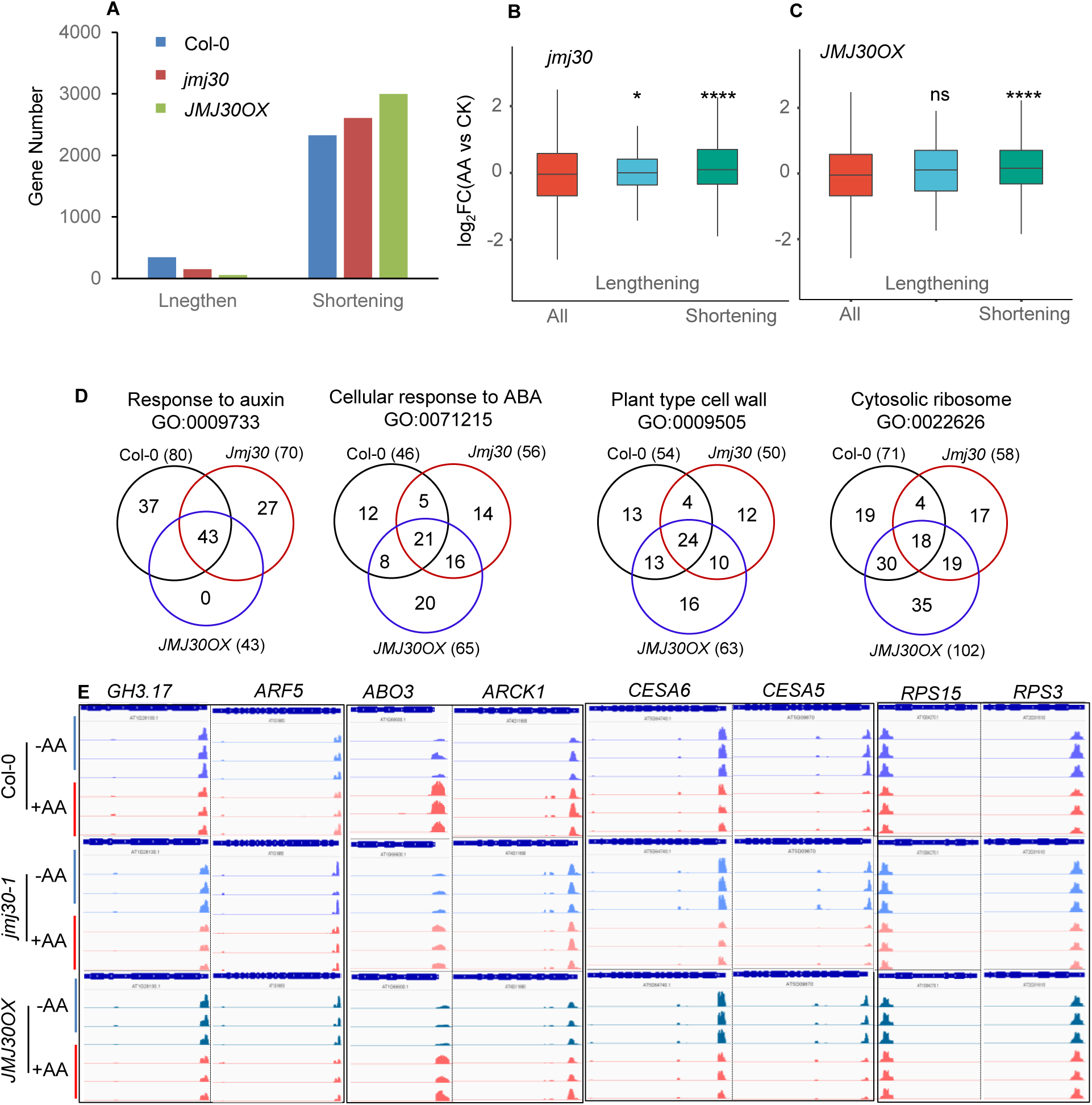
The histone demethylase JMJ30 regulates 3′ UTR shortening in response to AA. **A**, AA-induced alteration of 3′ UTR length. The number of genes with 3′ UTR shortening increases in both the *jmj30* mutant and *JMJ30* overexpression lines compared to wildtype seedlings in response to AA treatment. **B** and **C**, 3′ UTR shortening contributes to the higher relative levels of upregulated transcripts in response to AA treatment in both the *jmj30* mutant (**B**) and *JMJ30* overexpression lines (**C**). “All” refers to all expressed genes (raw read >0). **D**, Venn diagrams showing the extent of overlap between the number of genes whose transcripts undergo 3′ UTR shortening for the four indicated biological processes across the three genotypes. **E**, IGV illustration of 3′ UTR shortening for the transcripts of genes enriched in response to auxin, cellular response to abscisic acid stimulus, cytosolic ribosome, and plant-type cell wall. Read coverage is shown on the same scale for the indicated genes in wildtype Col-0, the *jmj30* mutant, and *JMJ30* overexpression seedlings. Note the differences in position and height of poly(A) signals between genotypes. See also a complete list of genes in Supplemental Data Set S3.

We visualized the PAS-seq read coverage in IGV for selected genes whose transcripts underwent 3′ UTR shortening in AA-treated and control samples. As shown in Figure 3E, we observed a switch in PAS position (as reflected by peak shape), together with a decrease in *CESA5* and *CESA6* PAS signal (as reflected by peak high) in all AA-treated seedlings. Moreover, we noticed an altered peak shape and height for *CESA6* PASs in control *jmj30-1* and *JMJ30OX* seedlings compared to Col-0. AA treatment changed the shape of *ABO3* and *ARCK1* PASs and resulted in a stronger induction of *ABO3* and *ARCK1* expression only in Col-0 and not in *jmj30-1* or *JMJ30OX*. Similar to *CESA5*, *CESA6*, *ABO3*, and *ARCK1*, the genes *GH3.17* and *ARF5* (*AUXIN RESPONSE FACTOR 5*) in “response to auxin” and *RPS15* (*CYTOSOLIC RIBOSOMAL PROTEIN S15*) and *RPS3* in “cytosolic ribosome” showed an alteration in the shape of PAS peaks in either AA-treated or control *jmj30-1* and *JMJ30OX* seedlings relative to Col-0 (Figure 3E). Together, these results indicate that JMJ30 is involved in the regulation of 3′ UTR shortening in response to mitochondrial stress.

### JMJ30 interacts with CPSF30 and loss of CPSF30 function confers mitochondrial stress and alters auxin and ABA responses

To further dissect JMJ30 function in the regulation of 3′ UTR processing upon mitochondrial stress, we identified JMJ30-interacting proteins using immunoprecipitation followed by mass spectrometry (IP-MS) analysis with plants overexpressing the *JMJ30-Myc* construct. A list of putative interacting proteins is provided in Supplemental Table S1. Among these proteins, we selected the cleavage and polyadenylation specificity factor CPSF30 for characterization. We established that CPSF30 interacts with JMJ30 by Co-IP, size exclusion chromatography (SEC), yeast two-hybrid (Y2H), and bimolecular fluorescence complementation (BiFC) assays (Figure 4, A-D). We conducted a SEC analysis to determine the size of the CPFS30-JMJ30 complex using Col-0 seedlings expressing both *YFP-CPSF30* (encoding a fusion between yellow fluorescent protein [YFP] and CPSF30) and *JMJ30-Myc* (encoding Myc-tagged JMJ30). We determined that CPSF30 and JMJ30 co-migrate with a larger apparent molecular mass than either protein alone, indicating that the two proteins are part of a complex in vivo (Figure 4B). The BiFC assay demonstrated that CPSF30 and JMJ30 interact in both the cytoplasm and the nucleus when their respective encoding constructs are co-infiltrated in *Nicotiana benthamiana* leaves (Figure 4D). We obtained similar results when testing the co-localization of YFP-CPSF30 and JMJ30-mCherry by confocal microscopy in *N. benthamiana* leaves (Figure 4E).

**Figure 4.**
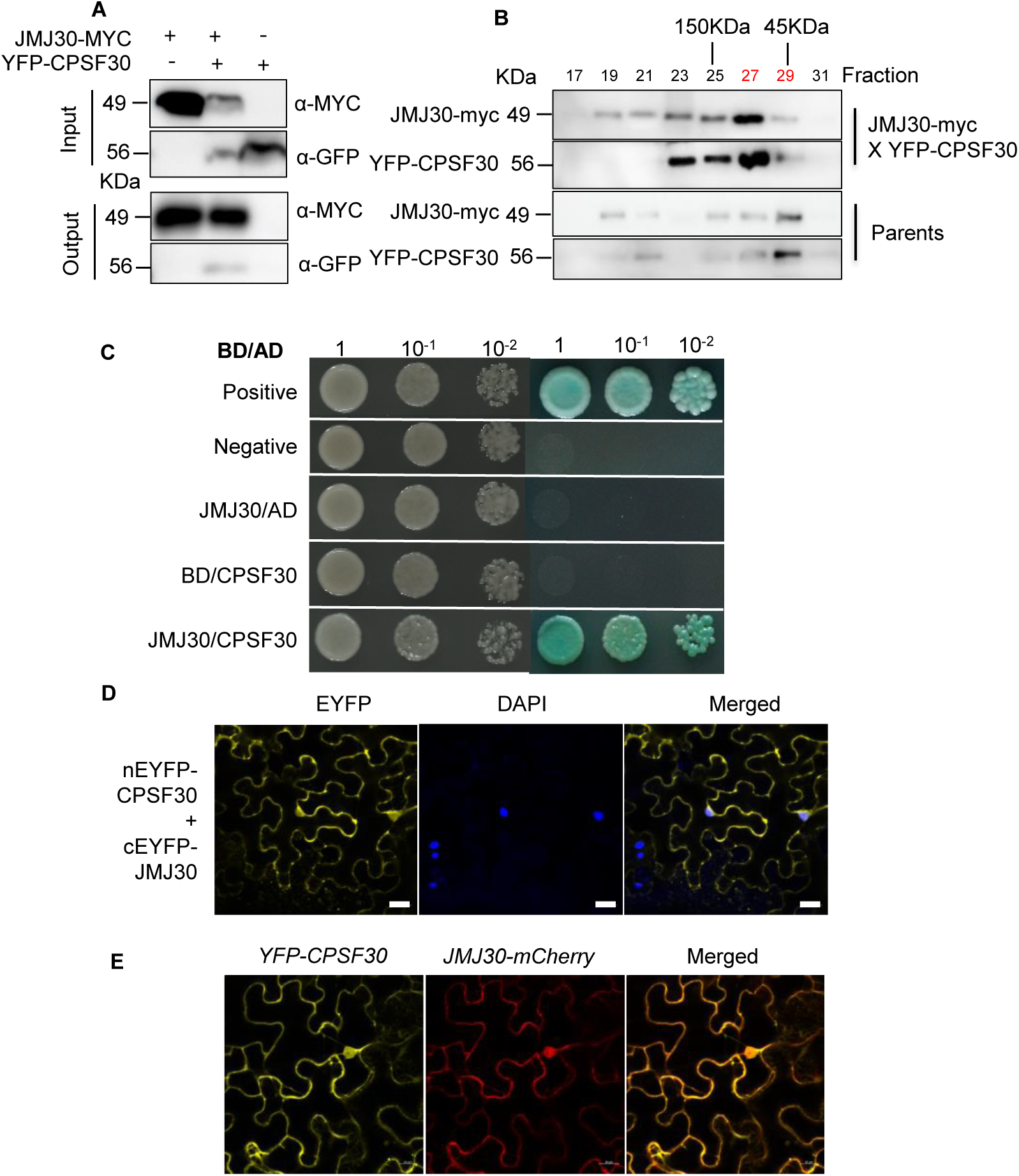
The histone demethylase JMJ30 interacts with the cleavage and polyadenylation specificity factor CPSF30. **A**, CPSF30 and JMJ30 interact by Co-IP. Co-IP assay showing that an anti-Myc antibody precipitates CPSF30-GFP when the encoding construct is co-infiltrated with *JMJ30-Myc* in *N. benthamiana* leaves, indicating that CPSF30-GFP and JMJ30-MYC are in the same protein complex. **B**, Analysis of the CPSF30-JMJ30 protein complex by size exclusion chromatography (SEC). CPSF30 and JMJ30 co-migrate in the same fraction with an approximate size of 110 kD, consistent with the formation of a protein complex. **C**, CPSF30 and JMJ30 interact in yeast. Yeast two-hybrid (Y2H) assay demonstrating that CPSF30 directly interacts with JMJ30. **D**, Detection of CPSF30 and JMJ30 interaction in the nucleus and cytoplasm by bimolecular fluorescence complementation (BiFC) assay. DAPI staining (blue) indicates the location of nuclei. **E**, Co-localization of CPSF30 and JMJ30 in the nucleus and cytosol of *N. benthamiana* cells co-infiltrated with *YFP*-*CPSF30* and *JMJ30-mCherry* constructs. Fluorescence was detected by confocal microscopy.

A function for CPSF30 in mitochondrion-to-nucleus communication has not been described. We thus obtained one *cpsf30* T-DNA insertional mutant, SALK_049389 (originally designated *oxidative stress tolerant6* [*oxt6-1*]), in which the T-DNA is inserted within the first exon (Supplemental Figure S5, A and B). The *CPSF30* locus is annotated as producing three splice variants; we determined that *CPSF30L* (At1g30460.1) and *CPSF30S* (At1g30460.3) transcript levels are less abundant in *oxt6-1* compared to Col-0 (Supplemental Figure S5, C and D). To explore a possible function for CPSF30 in mitochondrial stress response, we transformed the *oxt6-1* mutant with a construct harboring the *CPSF30* promoter driving the *CPSF30S* genomic coding region for complementation assays and overexpressed *CPSF30S* in Col-0. We observed a weaker inhibition of primary root growth in the *oxt6-1* mutant in the presence of AA compared to Col-0, indicating that loss of CPSF30 function results in a slight insensitivity to AA-induced growth inhibition (Figure 5, A and B). Notably, the growth of the primary root and the whole seedling of the *oxt6-1* mutant also appeared to respond differentially to auxin, ABA, and H_2_O_2_ treatment (Figure 5, A and B). The expression of the genomic fragment of *CPSF30S* driven by the *CPSF30* promoter complemented the phenotypes exhibited by the *oxt6-1* mutant in response to AA; the overexpression of *CPSF30S* produced a similar growth response to AA treatment as Col-0. In addition to AA responses, the *CPSF30S* complementation line and the overexpression line behaved similarly in response to ABA, auxin, and H_2_O_2_ treatment as the wildtype Col-0. These results clearly demonstrate that CPSF30S functions in AA-induced MRR and also plays a role in ABA and auxin signaling as well as in the oxidative stress response.

**Figure 5.**
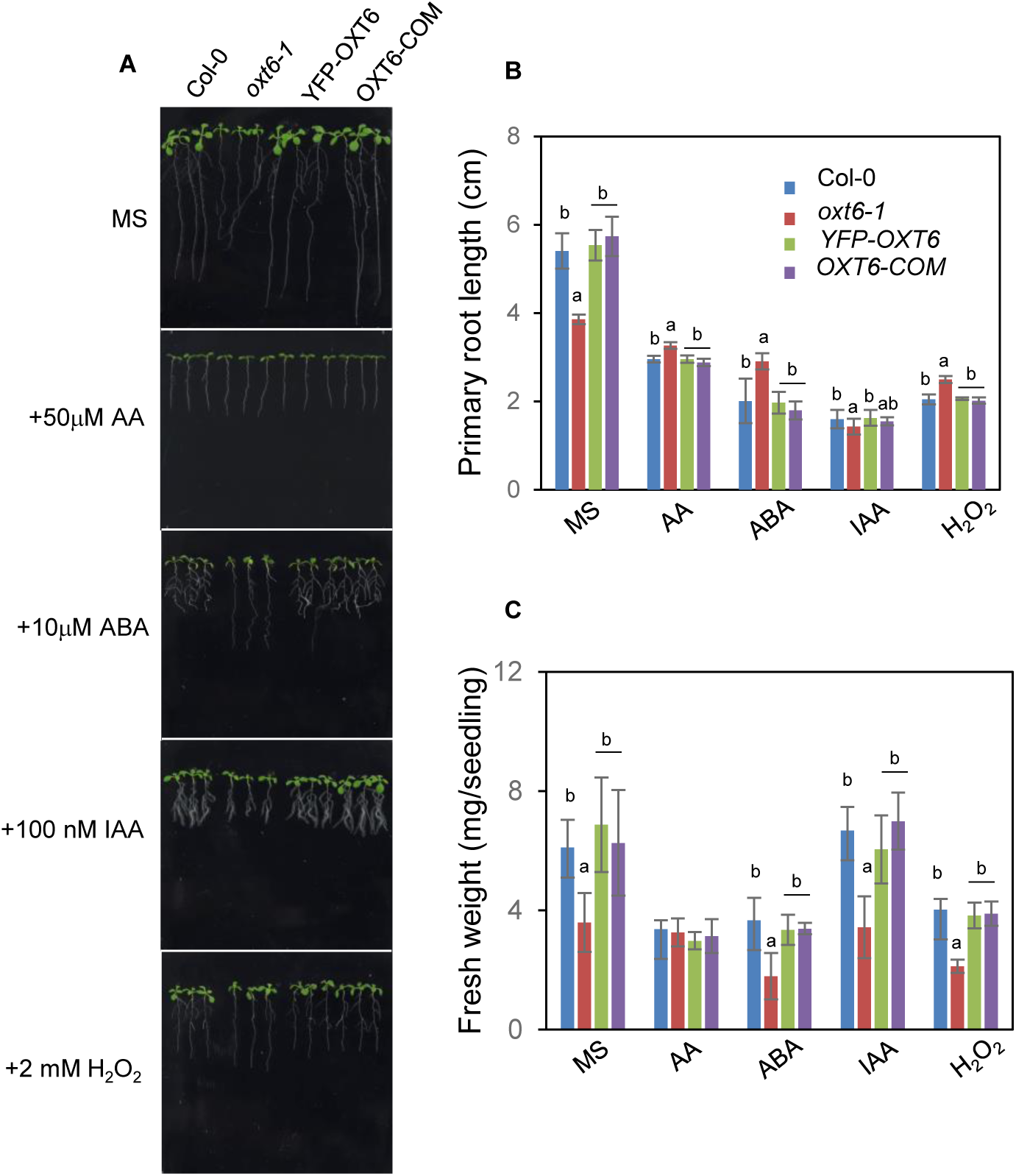
*CPSF30* expression complement *oxt6* mutant phenotype. **A**, Phenotype of the *oxt6* mutant and *CPSF30* transgenic lines. Seedlings from the *oxt6-1* mutant, an *oxt6-1* complemented line (harboring the *CPFS30* genome region driven by the *CPSF30* promoter, *CPSF30-COM*), Col-0 (wild type), and Col-0 expressing *YFP-CPSF30* from the CaMV 35S promoter (*YFP-CPSF30*) were grown on MS plates alone or containing 50 µM AA, 10 µM ABA, 100 nM IAA, or 2 mM H_2_O_2_. Photographs were taken 7 d after transfer to the treatment conditions. **B** and **C**, Mean root length (**B**) and fresh weight (**C**) of the seedlings shown in (**A**). Data are means ± SD (*n* = 3–9). One-way ANOVA was performed to determine significance. See also Supplemental Data Set S5 for statistical analysis.

### JMJ30 controls H3K27me3 levels at CPSF30-targeted loci

To address the biological significance of the JMJ30-CPSF30 interaction in the control of 3′ UTR processing, we generated the *jmj30-1 oxt6-1* double mutant by crossing and characterized its growth and 3′ UTR length in response to AA treatment. Growth responses of the *jmj30-1 oxt6-1* double mutant to AA, ABA, IAA, and H_2_O_2_ are similar to that of the *oxt6-1* single mutant (Supplemental Figure S6, A and B). We calculated the relative shortening index (RSI) for AA-treated and control seedlings for the wild type, the single mutants, and the double mutant (Figure 6A and B). We selected five genes from the 3′ UTR shortening dataset for each of the enriched GO terms auxin response, ABA response, cytosolic ribosome, and plant-type cell wall and assessed their 3′ UTR processing in *jmj30-1*, *oxt6-1*, and *jmj30-1 oxt6-1*. We used *ARF19* as a control, as it did not show 3′ UTR shortening (Supplemental Figure S7). For each GO process, the transcripts from the five respective genes, but not those of *ARF19*, had positive RSI values in the wild type, thus validating the PAS-seq analysis (Figure 6C). The RSI values of most transcripts whose genes were related to auxin response, cytosolic ribosome, and plant-type cell wall were significantly altered in the *oxt6-1* single mutant and the *jmj30-1 oxt6-1* double mutant relative to Col-0. Compared to the *oxt6-1* and *jmj30-1* single mutants, RSI values of several genes in the *jmj30-1 oxt6-1* double mutant followed the same trend as the wild type, suggesting that effects caused by the loss of CPSF30 function may be compensated for by enhancing *JMJ30* expression in the choice of PAS in response to AA treatment. Together, these results suggest that CPSF30 is required for AA-induced 3′ UTR shortening and that a potential regulation by JMJ30 and CPSF30 coaction may take place to modulate PAS usage in some loci in response to mitochondrial stress.

**Figure 6.**
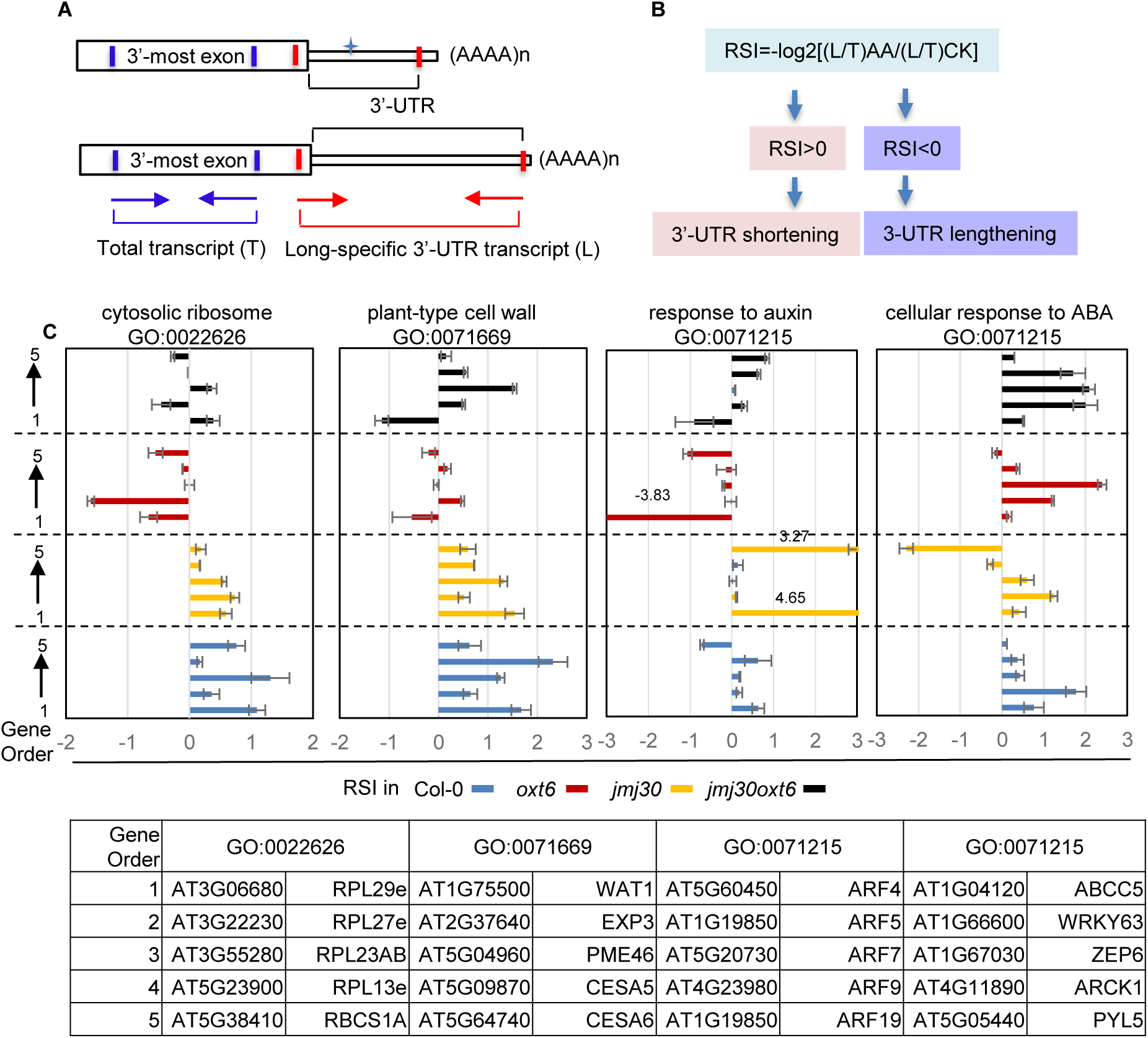
CPSF30 and JMJ30 regulate 3′ UTR shortening in response to AA. **A**, Schematic diagram of the primer sets used for the determination of relative shortening index (RSI). The primer pair for determining total transcript abundance (short +long) was designed within the 3′-most exon, and the pair used for determining the abundance of the long transcript was designed at the end of the 3′-most exon (forward) and the end of the 3′ UTR (reverse). Also see Supplemental Figure S7 for primer sets designed for each gene. **B**, Equation used to calculate RSI. RSI > 0 indicates 3′ UTR shortening; RSI < 0 indicates 3′ UTR lengthening in AA-treated samples (AA) vs control samples (CK) for each genotype. **C**, Comparison of RSI values between AA-treated and control samples. RSIs in Col-0, *oxt6-1*, *jmj30-1*, and *jmj30-1oxt6-1* are presented in blue, red, yellow, and black columns, respectively. GO terms are indicated at the top of the panel, and the genes examined in each GO category are listed below the panels.

Given that JMJ30 functions as a H3K27me3 demethylase and that CPSF30 is likely involved in auxin-regulated polyadenylation (Gan et al., 2014; Hong et al., 2018), we performed ChIP-PCR to check the H3K27me3 status at the *ARF5*, *ARF7*, and *CESA6* loci in *jmj30-1*, *oxt6-1*, *jmj30-1 oxt6-1*, and *oxt6-COM* compared with the wild type, using *ACT2* as a negative control. We targeted five regions across the entire gene: the promoter, near the ATG, within the last coding exon, and near the first (pA1) or second (pA2) poly(A) site within the 3′ UTR (Supplemental Figure S8A). We observed that the H3K27me3 levels at *CESA6*, *ARF5*, and *ARF7*, but not at *ACT2*, are all significantly high in *oxt6-1*, but relatively low in *jmj30-1* compared to the wild type (Figure 7, A-D). Interestingly, the H3K27me3 level in the region near the ATG in *ARF5* and *ARF7* did not significantly increase relative to the wild type, suggesting that the consequences of the loss CPSF30 function on H3K27me3 are biased toward the last coding exon and the 3′ UTR. *CESA6* exhibited a similar pattern, although we also detected high levels of H3K27me3 in the region near the ATG. Importantly, H3K27me3 levels were comparable in the wild type and *oxt6-COM* at three loci, demonstrating a functional complementation of the *oxt6-1* mutant by *CPSF30S* at the molecular level. As we observed similar H3K27me3 levels in the wild type and *jmj30-1 oxt6-1*, the suppression of high H3K27me3 levels by JMJ30 in the *oxt6-1* background suggested that enhancing *JMJ30* expression reduces H3K27me3 levels at CPSF30 targeted loci.

**Figure 7.**
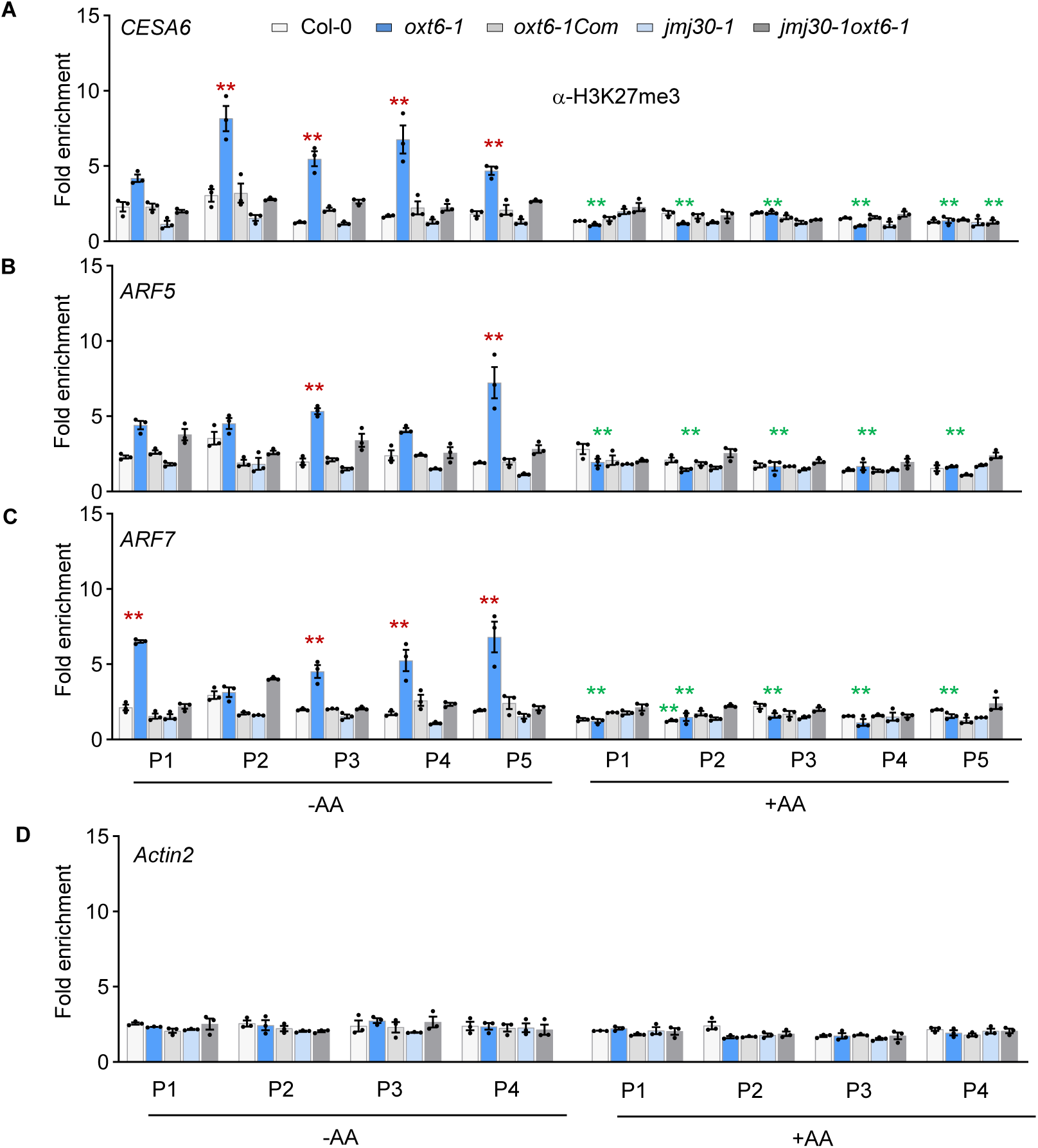
JMJ30 reduces H3K27me3 enrichment at CPSF30 targeted loci. **A-D**, ChIP-qPCR analysis of H3K27me3 status at the *CESA6* (**A**), *ARF5* (**B**), *ARF7* (**C**), and *ACT2* (**D**) loci in the Col-0, *oxt6-1*, *jmj30-1*, and *oxt6-1 jmj30-1* backgrounds. The seedlings were treated with (+AA) or without (– AA) 50 μM AA for 5 h. H3K27me3-specific fold enrichment (FE) of each examined region was normalized to the non-specific fold enrichment values obtained with an anti-GFP antibody. Also see the detailed examined regions of all loci in Supplemental Figure S8. Student’s *t*-test was performed to determine significance. Significant changes in H3K27me3-specific fold enrichment were defined as: FE > 2, with \**P* < 0.05, \*\**P* < 0.01. Data are means ± SD (*n* = 3). Red stars indicate significant increases in H3K27me3 levels at a given locus in the *oxt6-1* mutant vs wildtype Col-0; green stars indicate significant decreases in H3K27me3 levels at a given locus for each AA-treated sample vs its corresponding control sample. See also Supplemental Data Set S6 for primer sets designed for each gene.

We detected only slight decrease in the levels of H3K27me3 in some region of the *CESA6*, *ARF5*, or *ARF7* loci in AA-treated samples compared to control samples, with only AA-treated *oxt6-1* seedlings exhibiting a 2- to 6-fold reduction in H3K27me3 levels relative to control seedlings (Supplemental Data Set S5). To assess whether JMJ30 directly controls H3K27me3 levels at *CPSF30*, we examined the H3K27me3 status in different regions of *CPSF30S* and *CPSFL* in the wild type and *jmj30-1*. Six regions include the promoter (P1) and near the ATG (P2) that is shared by *CPSF30S* and *CPSF30L*, the last coding exon and the 3′ UTR that are unique to *CPSF30S* and *CPSF30L*, designed as P3-S and P3-L, P4-S and P4-L, respectively (Supplemental Figure S8B). We detected no significant differences for most regions of *CPSF30S* or *CPSF30L* in *jmj30-1* compared to wild type with or without AA treatment, indicating that JMJ30 has no major effects on H3K27me3 levels at *CPSF30*.

Together, these results suggest that CPFS30 is required for JMJ30 to remove the H3K27me3 mark at CPSF30 target loci. JMJ30 and CPSF30 have common targets, and the action of JMJ30 and CPSF30 may be interdependent.

### The JMJ30-CPSF30 module mediates mitochondrion-directed transcriptional regulation of the phytohormone pathway and cell wall biogenesis

Transcripts whose 3′ UTR underwent shortening in response to AA treatment were significantly enriched in the GO terms “response to auxin” and “cell wall biogenesis”, suggesting that these biological processes are involved in the co-transcriptional response to mitochondrial stress. We thus asked whether the CPSF30-JMJ30 module regulated the transcriptional response of these biological processes to changes in mitochondrial functional state by performing an RNA-seq analysis of AA-treated and control wildtype, *jmj30-1*, *oxt6-1*, and *jmj30-1 oxt6-1* seedlings. A comparison of transcript levels between AA-treated and control seedlings identified 2,710 (Col-0), 2,558 (*jmj30-1*), 3,477 (*oxt6-1*), and 2,797 (*jmj30-1 oxt6-1*) DEGs (Supplemental Data Set S4). Among them, we detected 1,135 AA-induced common DEGs across all genotypes (adjusted *P*-value < 0.05) (Figure 8A). Among the top biological processes enriched in DEGs, we focused on phytohormone response (Figure 8B), cell surface receptor signaling, cell wall organization, and cell growth (Figure 8C). We observed that ABA-, ethylene-, JA-, and SA-responsive genes and phytohormone biosynthetic genes are all induced by AA treatment, with a subset of auxin-responsive genes downregulated upon AA treatment. Interestingly, these phytohormone-responsive genes tended to be strongly repressed in the *oxt6-1* mutant relative to the wild type (Figure 8B and D) but were highly upregulated in AA-treated *oxt6-1* seedlings, highlighting a prominent transcriptional signature among AA-treated samples. Another notable transcriptional signature was the much greater transcriptional downregulation of components of cell surface receptor signaling, cell wall organization, and cell growth in the *jmj30-1* mutant compared to other AA-treated samples (Figure 8C and E, Supplemental Data Set S5). For example, the cellulose synthase genes *CESA1*, *CESA2*, *CESA4*, *CESA5*, and *CESA6*, involved in cell wall biogenesis and assembly, were all significantly repressed in AA-treated *jmj30-1* mutant seedlings compared to AA-treated Col-0 (Figure 8E, Supplemental Data Set S5). These results suggest that the JMJ30-CPSF30 module controls the transcription of genes involved in phytohormone pathways and cell wall biosynthesis to sense mitochondrial functional status.

**Figure 8.**
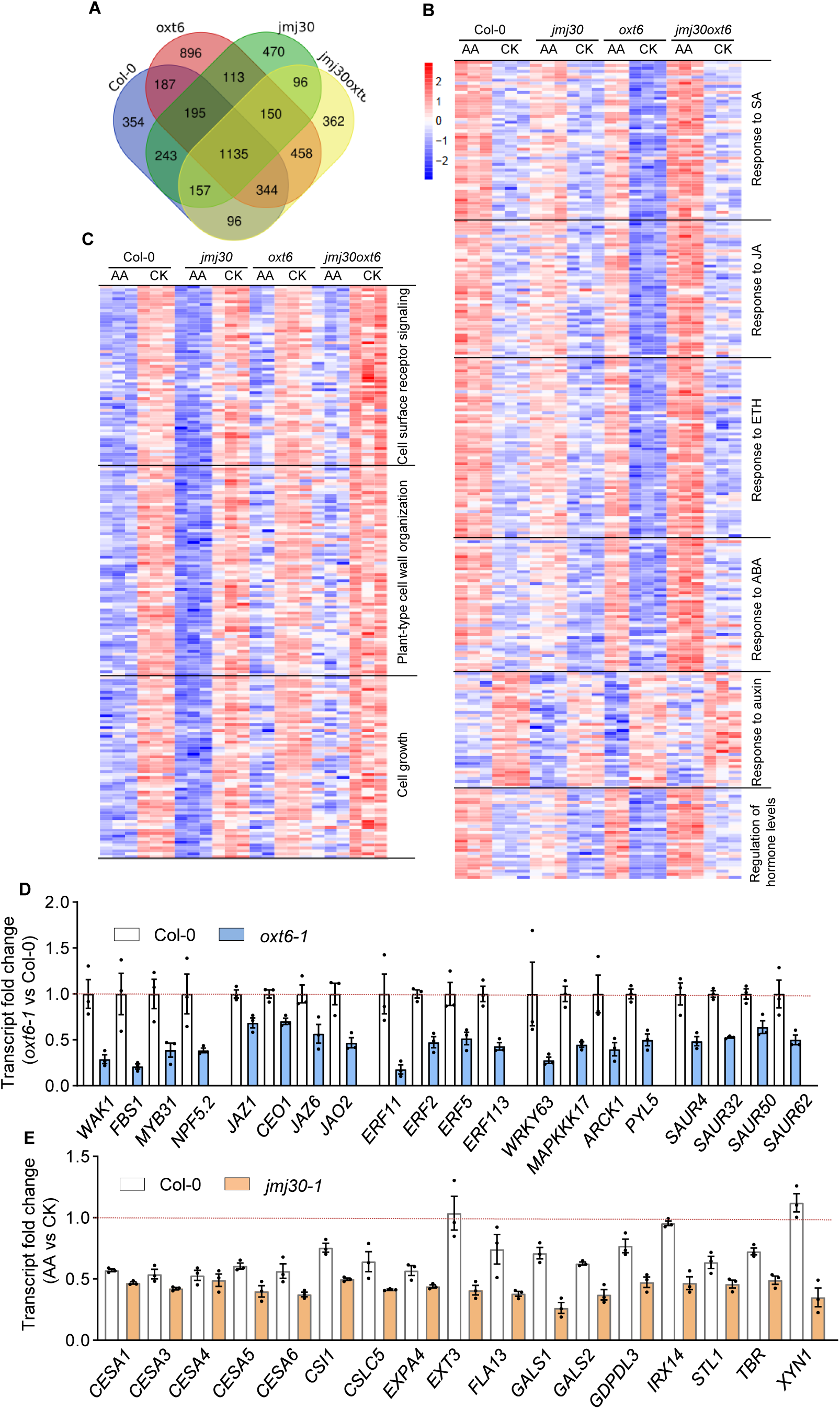
The expression of phytohormone-responsive genes is highly induced in *oxt6-1* and *jmj30-1 oxt6-1* mutants, while that of growth-related genes is strongly repressed in *jmj30-1* upon AA-induced mitochondrial stress. **A**, Venn diagram showing the extent of overlap between differentially expressed genes (DEGs) identified in four AA-treated samples relative to their controls. **B**, Heatmap representation of expression levels from upregulated and downregulated phytohormone-responsive genes among 1,135 DEGs common to all genotypes. The associated enriched GO terms “response to salicylic acid” (GO:0009751), “response to jasmonic acid” (GO:0009753), “response to ethylene” (GO:0009723), “cellular response to abscisic acid stimulus” (GO:0071215), “response to auxin” (GO:0009733), and “regulation of hormone levels” (GO:0010817) are indicated to the right of or below the heatmap. **C**, Heatmap representation of expression levels from downregulated growth-related genes among the 1,135 common DEGs. The associated enriched GO terms “cell surface receptor signaling” (GO:0007166 and GO:0007167), “plant-type cell wall organization” (GO:0071669), and “cell growth” (GO:0016049) are indicated to the right of the heatmap. **D** and **E**, Relative transcript levels for selected plant hormone–responsive genes in Col-0 and the *oxt6-1* mutant (**D**) or the *jmj30-1* mutant (**E**), as determined by RNA-seq and reported as fold-change in *oxt6-1* control samples vs wildtype control samples (**D**) or fold-change of AA-treated vs control samples (**E**). Selected phytohormone-responsive genes and cell wall–related genes were all significantly downregulated in *oxt6-1* and in AA-treated *jmj30-1* mutants compared to the wild type. Data are means ± SD (*n* = 3). See also Supplemental Data Set S4 for details.

One possible mechanism by which mitochondria might affect the activity of plant hormone pathways is though the regulation of phytohormone levels. We thus measured plant hormone levels in *oxt6-1* and wildtype seedlings upon AA treatment. The assay used in this study allowed the simultaneous measurement of main phytohormones and their derivatives in each sample. We detected no significant changes for ABA, SA, or JA contents in AA-treated wildtype or *oxt6-1* seedlings compared to control seedlings, while the GA precursor GA_24_ accumulated to slightly lower levels in AA-treated wildtype and *oxt6-1* seedlings (Figure 9A and B). Compared to the wild type, ABA and 1-aminocyclopropane 1-carboxylic acid (ACC) contents were higher in *oxt6-1* with or without AA treatment. We also observed a slight but significant reduction in ACC contents for AA-treated Col-0 (Figure 9B). For cytokinins, we observed that isopentyladenine (iP, active cytokinin), isopentenyl riboside (iPR), and trans-zeatin-riboside (tZR) (inactive cytokinin derivatives) accumulate to significantly lower levels, while the contents of cis-Zeatin riboside (cZR, inactive cytokinin derivative) and 6-benzylaminopurine (BAP, active cytokinin) significantly increased in the AA-treated wild type. In *oxt6-1*, similar to the wild type, IP, IPR, and BAP levels decreased, while those of tZR, but not cZR, strongly increased upon AA treatment (Figure 9C). These results demonstrate that AA treatment has no significant effects on ABA, SA, JA, or potential ethylene biosynthesis, but does to some extent affect cytokinin metabolism. CPSF30 showed negative regulatory roles in ABA and ACC biosynthesis.

**Figure 9.**
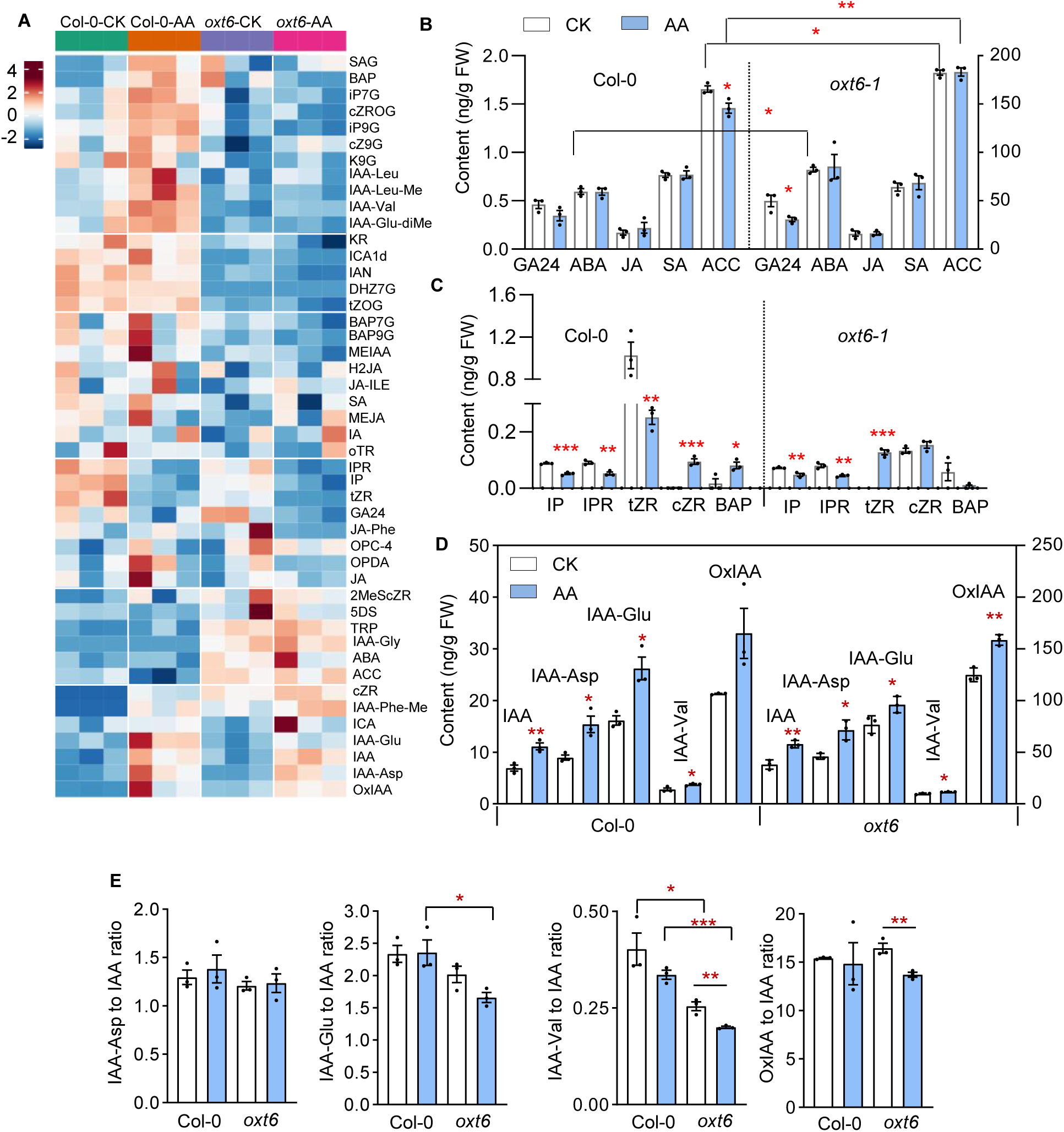
Response of phytohormone levels to AA-induced mitochondrial stress. **A**, Heatmap representation of phytohormone contents in AA-treated and control *oxt6-1* and wildtype seedlings. IAA, ABA, cytokinin, ACC, JA, SA, and their derivatives are indicated to the right of the panel, and sample groups are indicated on the top of panel. A scale bar is shown to the left. **B**, Mean levels for ABA, JA, SA, ACC, and GA_24_ in *oxt6-1* and Col-0. Note the lower levels of ACC and GA_24_ in AA-treated Col-0 and *oxt6-1*, respectively. GA_24_ and ABA contents are plotted along the left y*-*axis; JA, SA, and ACC contents are plotted along the right y*-*axis. **C**, Mean levels of cytokinin and their derivatives in *oxt6-1* and Col-0 upon AA treatment. **D**, Mean levels of IAA and their derivatives in *oxt6-1* and Col-0 upon AA treatment. Only oxIAA contents are plotted along the right y*-*axis. **E**, Mean ratio between IAA derivatives and free IAA in *oxt6-1* and Col-0 upon AA treatment. Data are means ± SD (*n* = 3). Student’s *t*-test was performed to determine significance. \**P* < 0.05, \*\**P* < 0.01. See also Supplemental Data Set S5 for statistical analysis.

Interestingly, the contents for IAA and its amino acid conjugates, IAA-Asp, IAA-Glu, and IAA-Val, as well as its oxidized form OxIAA, all significantly increased in both AA-treated wildtype and *oxt6-1* seedlings compared to controls, indicating that IAA production is enhanced upon AA treatment (Figure 9, A and D, Supplemental Data Set S5). We calculated the ratio between IAA-aa (amino acid conjugates) or OxIAA and free IAA in all genotypes. These ratios were significantly lower in *oxt6-1* compared to the wild type, suggesting that free IAA accumulates to relatively higher levels in *oxt6-1* than in the wild type upon AA treatment (Figure 9E). We also observed that Trp content was slightly higher in AA-treated samples but significantly higher in *oxt6-1* than in the wild type (Figure 10A). We also examined the expression levels of genes involved in Trp-dependent IAA biosynthesis, which revealed that *YUCCA5* (*YUC5*) and *YUC8* expression levels significantly increase in AA-treated *oxt6-1*, while other *YUC*s were slightly increased or remained unchanged in AA-treated wildtype and *oxt6-1* seedlings compared to controls (Figure 10B). Compared to the wildtype, *oxt6-1* showed high *YUC3* and *YUC4* transcript levels, but low *YUC5* and *YUC6* levels (Figure 10C). *AMIDASE 1* (*AMI1*) transcript levels were also slightly induced by AA in the wildtype, while *oxt6-1* showed a high basal level of *AMI1* transcripts compared to the wildtype (Figure 10D). The increase of AA-induced IAA-aa may be attributed to both AA-induced IAA production and AA-induced upregulation of IAA conjugate acyl amido synthetases from the GH3 family that conjugates IAA to amino acids in both AA-treated wildtype and *oxt6-1* seedlings (Figure 10E). The transcription of all *GH3* family members was activated in AA-treated *oxt6-1*, although most *GH3* transcript levels were comparable across control samples (Figure 10F). These results suggest that AA induces auxin production likely via a Trp-dependent pathway. Free IAA levels were relatively high in AA-treated and control *oxt6-1* seedlings compared to the wild type.

**Figure 10.**
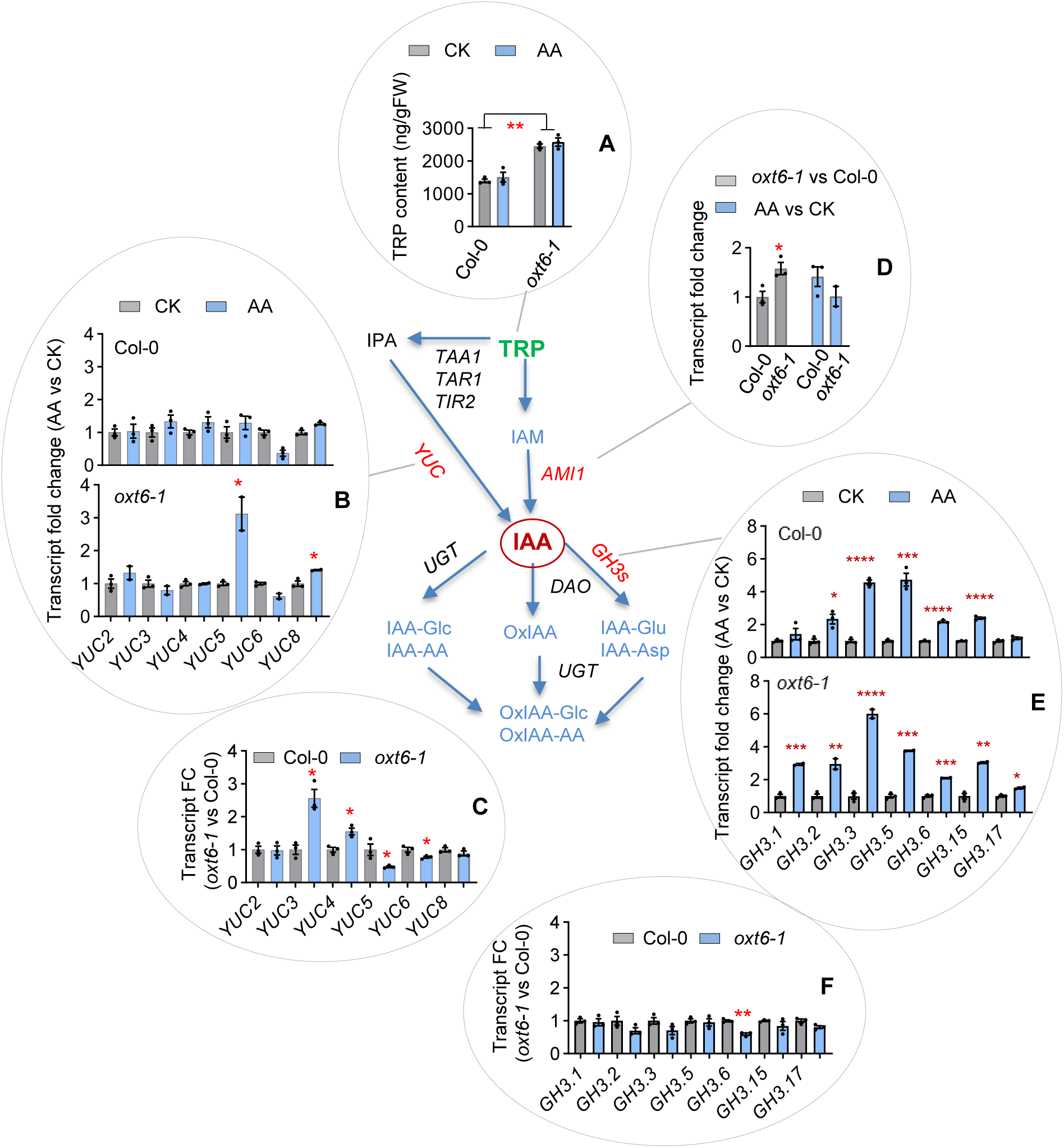
Regulation of genes in Trp-dependent auxin biosynthesis in *oxt6-1* and Col-0 upon AA-induced stress. **A** simplified diagram of Trp-dependent auxin biosynthesis pathway is shown in the center. Mean contents of the IAA precursor Trp and transcript levels in the pathway in *oxt6-1* and Col-0, or AA-treated samples vs control samples, as indicated. A, Trp contents are significantly higher in *oxt6-1* than in Col-0 with or without AA treatment. **B**, The transcript levels of most *YUC* family members are induced slightly by AA in Col-0, but *YUC5* and *YUC8* transcript levels significantly increase upon AA treatment in *oxt6-1*. **C**, *YUC3* and *YUC4* transcript levels are significantly higher but *YUC5* and *YUC6* levels are significantly lower in *oxt6-1* compared to Col-0. **D**, *AMI1* expression levels are high in *oxt6-1* compared to Col-0. AA induces no significant changes of *AMI1* expression. **E**, The expression of *GH3* family members is significantly induced upon AA treatment in both *oxt6-1* and Col-0. **F**, No significant differences are detected for the transcript levels of most *GH3* family members except for *GH3.6* between *oxt6-1* and Col-0. Data are means ± SD (*n* = 3). Student’s *t*-test was performed to determine significance. \**P* < 0.05, \*\**P* < 0.01. See also Supplemental Data Set S5 for statistical analysis.

Taken together, we conclude that AA-induced IAA production likely acts via a Trp-dependent pathway and that free IAA levels are relatively high in the *oxt6-1* mutant. The activity of phytohormone pathways may not be directly determined by phytohormone levels during mitochondrial stress. Namely, the enhanced expression of phytohormone-responsive genes may not necessarily be correlated with an increase in phytohormone levels because AA has no effects on their contents, or show the opposite of effect.

### Perturbation of auxin biosynthesis increases the sensitivity to growth inhibition, but the repression of cell wall biogenesis benefits plant fitness under mitochondrial stress

To determine the biological relevance of the regulation of cellular processes mediated by the JMJ30-CPSF30 module under mitochondrial stress, we examined the growth phenotype of mutants defective in auxin biosynthesis or transport, as well as cell wall biogenesis–related mutants under AA-induced mitochondrial stress. We obtained the *yuc* mutant *yuc9* and the *yuc1 yuc2 yuc4 yuc6* quadruple mutant, the *pin* mutants *pin1-11* and *pin2*, and the cell wall mutants *cesa3*, *cesa6*, *mur1*, and *mur4* and examined their growth under AA treatment. Notably, the *yuc1 yuc2 yuc4 yuc6* quadruple mutant showed a more severe AA-induced growth inhibition than the wild type or the *yuc9* single mutant (Figure 11, A and B). The growth of both *pin1-11* and *pin2* was also more sensitive to AA treatment than the wild type (Figure 11, A and B). CESA3 and CESA6 function in primary cell wall formation, while MUR1 and MUR4 play roles in cell wall modification. Their corresponding cell wall mutants were all less affected by AA-induced growth inhibition and were partially insensitive to AA treatment for primary root and whole seedling growth (Figure 11, C and D). Together, these results suggest that disturbing auxin biosynthesis reduces cellular tolerance to mitochondrial stress, whereas reducing cell wall formation confers greater plant fitness while mitochondrial function is disrupted.

**Figure 11.**
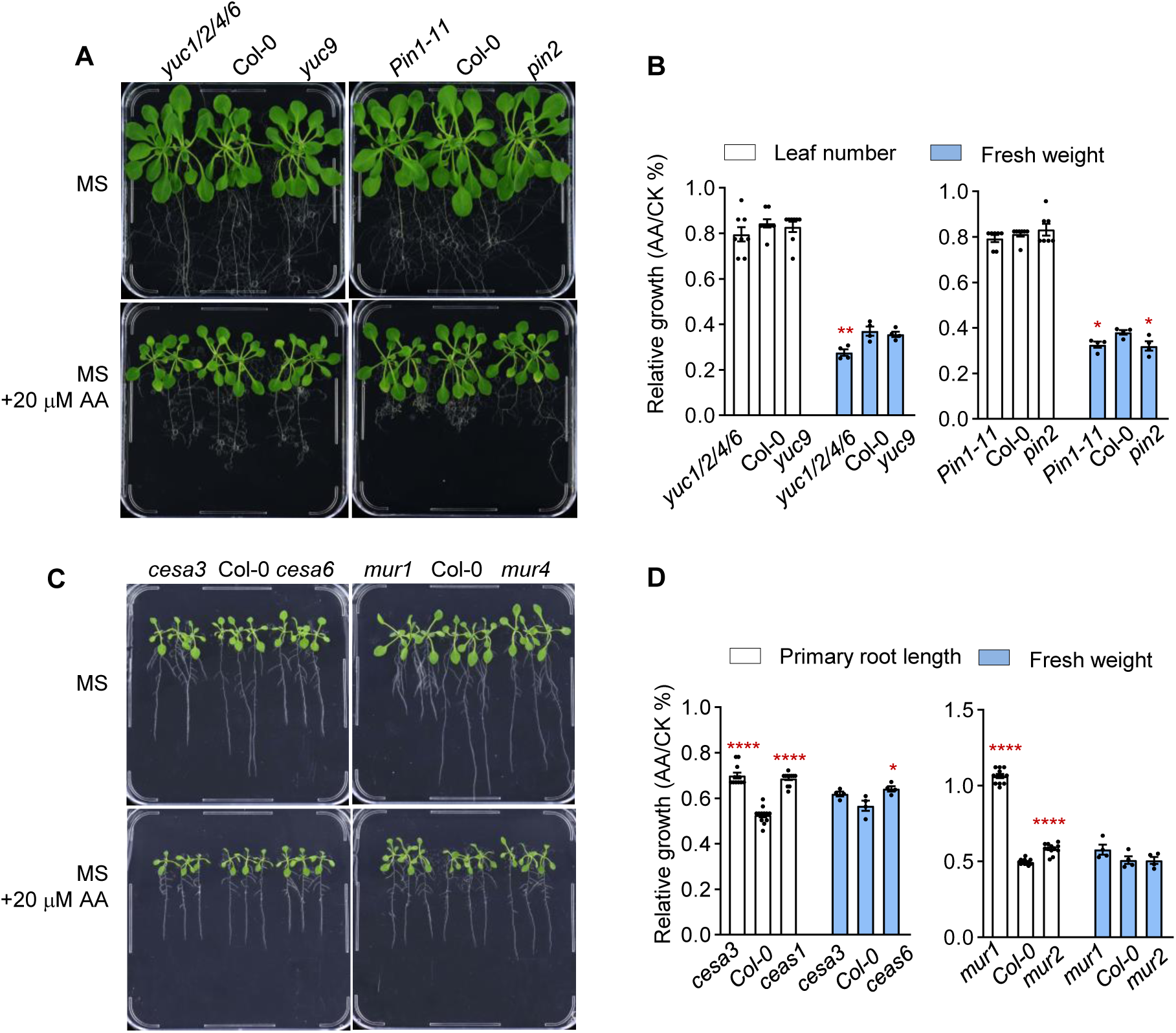
Growth assessment of auxin and cell wall mutants under AA-induced mitochondrial stress. **A**, Seedling growth of auxin biosynthetic mutants and Col-0 upon AA treatment. Upper panel, seedlings grown on MS medium; lower panel, seedlings grown on MS medium supplemented with 20 µM AA. Photographs were taken around 2 weeks after 5-d-old seedlings were transferred to MS medium or MS medium containing 20 µM AA. **B**, Summary of growth parameters of the seedlings shown in (**A**). Leaf number and fresh weight of AA-treated seedlings were normalized to values from control seedlings. **C**, Seedling growth of cell wall mutants and Col-0 upon AA treatment. Upper panel, seedlings grown on MS medium; lower panel, seedlings grown on MS medium supplemented with 20 µM AA. Photographs were taken about 1 week after 5-d-old seedlings were transferred to MS medium or MS medium containing 20 µM AA. **D**, Summary of growth parameters of the seedlings shown in (**C**). Primary root length and fresh weight of AA-treated seedlings were normalized to values from control seedlings. Student’s *t*-test was performed to determine significance. \**P* < 0.05, \**P* < 0.01. Data are means ± SD (*n* = 3). See also Supplemental Data Set S5 for statistical analysis.

## Discussion

To establish the potential function of mRNA 3′-end processing and epigenetic regulation in mitochondrion-to-nucleus communication, we examined global AA-induced APA events in wildtype seedlings and gain-of-function mutants in the histone H3K27me3 demethylase JMJ30. Our findings support a model whereby mitochondrion functional state regulates nuclear alternative mRNA polyadenylation, with an influence on nuclear gene expression, thus connecting mitochondrial retrograde signaling with alternative nuclear mRNA polyadenylation. Although the nature of the retrograde signals that are relayed from mitochondria to the nucleus to modulate APA are unclear, we demonstrated the participation of the epigenetic effector, the histone demethylase JMJ30, and the polyadenylation specificity factor CPSF30 in this process. Moreover, our study identifies auxin responses and cell wall biogenesis as the main target pathways of AA-induced 3′ UTR shortening that shape growth and fitness. Thus, our results clearly indicate that APA and epigenetic modifications enable quick cellular responses to mitochondrial stress while producing multiple RNA variants in the nucleus and changing translation efficiency in the cytoplasm for growth and adaptation.

We showed that about 60% of Arabidopsis genes have at least two PASs (Supplemental Figure S1C), which is similar to the number reported in previous studies (Hong et al., 2018; Lin et al., 2020b). We also demonstrated that the 3′ UTR length of most transcripts undergoes dynamic changes upon AA-induced mitochondrial stress, with a preference for a proximal PAS. Although a precise mechanism(s) by which AA-induced mitochondrial stress leads to 3′ UTR shortening of the selected transcripts is unknown, we established that enhancing *JMJ30* expression levels increases the number of transcripts whose 3′ UTR undergoes shortening (Figure 3A). Most known 3′-end processing factors do not alter their gene expression in the AA-treated wild type, but some were upregulated or downregulated upon AA treatment in the *jmj30-1* mutant (Supplemental Figure S9), suggesting that AA-induced 3′ UTR shortening occurs by the co-action of multiple factors, presumably including different JMJ30 interacting APA factors that have not been characterized in this study (Supplemental Table S1). However, the physical interaction between JMJ30 and CPSF30 offered a potential mechanism for the epigenetic regulation of AA-induced APA through 3′-end processing. Although CPSF30 is a core subunit of the CPSF, its precise roles in polyadenylation remain unclear. Previous studies have indicated that CPSF30 interacts with many other CPSF subunits as well as components of the polyadenylation complex (Hunt et al., 2008; Zhao et al., 2009), making it a potential mediator of regulated APA in Arabidopsis, in addition to its direct RNA binding and endonuclease activity in pre-mRNA processing prior to poly(A) addition (Addepalli and Hunt, 2007; Thomas et al., 2012). CPSF30 has been implicated in oxidative stress response, auxin-mediated root development, flowering time, and nitrate metabolism (Zhang et al., 2008; Li et al., 2017; Hong et al., 2018; Yu et al., 2019a; Hou et al., 2021). Arabidopsis *CPSF30* has three distinct transcripts; the shorter transcript encodes a 28-kD protein (CPSF30S) and the longer transcript encodes a 70-kD protein (CPSF30L). CPSF30L has a unique YT521-B homology (YTH) domain that specifically recognizes N^6^-methyladenosine (m^6^A), conferring a function as an m^6^A reader and a regulator of APA for nitrate signaling–related genes (Zhang et al., 2008; Hou et al., 2021; Song et al., 2021). Notably, neither *CPSF30S* nor *CPSF30L* was a direct target of JMJ30, as we detected no significant changes in H3K27me3 levels across the *CPSF30S* or *CPSF30L* gene in the *jmj30-1* mutant (Supplemental Figure S7B). Consistently, *JMJ30* expression levels and AA treatment had no significant effects on total *CPSF30* transcript levels (Supplemental Figure S10, A and B), indicating that the alteration of 3′-end processing in the *jmj30-1* mutant is not related to *CPSF30* expression. However, the loss of CPSF30 function resulted in the significant increase of H3K27me3 levels at loci involved in auxin signaling and cell wall biogenesis, which was fully complemented by the expression of *CPSF30S* driven by the *CPSF30* promoter (Figure 7). This result suggests that m^6^A-regulated APA may not be involved in this process. We detected an increased H3K27me3 status over the 3′ end before and/or near the PAS in the *oxt6-1* mutant, as well as in the promoter, and near the ATG of *CESA6*, suggesting the potential influence of H3K27me3 levels in transcription, in addition to 3′-end processing. This hypothesis is supported by RNA-seq analysis, which revealed the very low transcript levels of most phytohormone-responsive genes in the *oxt6-1* mutant, but their similar levels in the *jmj30-1 oxt6-1* double mutant compared to the wildtype (Figure 8). These results suggest that CPSF30 is required for JMJ30 function at CPSF30-target loci and that raising *JMJ30* expression partially compensates for the loss of CPFS30 at the molecular level. Therefore, we propose that CPSF30 facilitates the recruitment of JMJ30 at CPSF30-target loci. CPSF30 and JMJ30 impart a transcriptional and co-transcriptional control on their common target loci involved in phytohormone response and cell wall biogenesis interdependently. JMJ30 activity reduces H3K27me3 levels and ensures an open chromatin structure that presumably enhances the recruitment of the 3′-end processing complex at proximal PAS, where JMJ30 and/or CPSF30 may also recruit other components of the 3′-end processing complex through direct or indirect physical interactions. Because *jmj30-1 oxt6-1* double mutant shows similar growth phenotype to *oxt6-1* single mutant under mitochondrial stress, we postulate that the recruitment of other components for 3′-end processing is CPSF30-dependent.

APA function in global control of gene expression is unclear in plants. Our results highlighted the rather weak association between AA-induced 3′ UTR shortening and either AA-induced downregulated or upregulated genes in the wild type. However, we observed a rise in the extent of correlation between AA-induced 3′ UTR shortening and AA-induced upregulated genes, but not with AA-induced downregulated genes with higher *JMJ30* expression, indicative of a regulatory role for JMJ30-mediated APA in the control of global gene expression. The influence of APA on gene expression is also reflected by the changes in the transcriptome observed in the *oxt6-1* mutant and the *jmj30-1 oxt6-1* double mutant relative to the wild type, suggesting a function for CPSF30 in the regulation of gene transcription via coaction with JMJ30. In addition, the analysis of 3′ UTR shortening and its relation to TE allowed the identification of biological processes that are regulated by 3′ UTR–controlled translation efficiency. We determined that 3′ UTR shortening, but not 3′ UTR lengthening, significantly contributes to translational regulation, which is similar to studies in mammalian cells and mice (*Mus musculus*) tissues that showed that the kinase mammalian Target of Rapamycin (mTOR) selectively regulates protein biosynthesis by modulating 3′ UTR length of mRNA (Chang et al., 2015). Interestingly, 3′ UTR–shortened transcripts with high TE were enriched in GO terms related to the cell wall, and those with low TE were enriched for GO terms associated with cell growth, whereas 3′ UTR– shortened transcripts among auxin response and cytosolic ribosome genes showed either high TE or low TE (Figure 2, Supplemental Figure S2). The TE and TS cross-analysis revealed that translation counteracts transcription, for example, for the genes involved in cell wall biogenesis and assembly, whereas they acted synergistically for ABA-responsive genes. For cytosolic ribosome genes with a shortened 3′ UTR upon AA treatment, TE may be the main determinant to their protein production rate. Together, these results clearly demonstrate that 3′ UTR shortening contributing to translation regulation is a regulatory mechanism governing mitochondrion-to-nucleus communication.

GO analysis of AA-induced genes whose transcripts underwent 3′ UTR shortening revealed that auxin responses and cell wall biogenesis are pathways targeted by APA that potentially mediate mitochondrion-to-nucleus communication. Given the important roles of mitochondria in plant growth and stress responses, it is not surprising that mitochondria would control growth and stress response through modulating the activity of phytohormone signaling and cell wall integrity. Auxin has been considered a major player in mitochondrion retrograde signaling; however, the cause-and-effect relationship between them was not clear (Berkowitz et al., 2016; Welchen et al., 2021). Our study clearly demonstrates that AA treatment increases IAA accumulation, presumably via a Trp-dependent pathway. Evidence supporting this claim includes 1) an increase in IAA levels in both AA-treated wild type and the *oxt6-1* mutant; 2) a slight increase in the expression of the *YUC* family members *YUC3*, *YUC4*, *YUC5*, and *YUC8* in the wild type, and the stronger induction of *YUC5* and *YUC8* in AA-treated *oxt6-1* seedlings; 3) the upregulation of the *GH3* family members that encode the enzymes that catalyze the conjugation of auxin to amino in both AA-treated wildtype and *oxt6-1* mutant seedlings; and 4) the high Trp levels, *AMI1*, *YUC3*, and *YUC4* transcript levels, and AA-induced free IAA contents in the *oxt6-1* mutant compared to the wild type. Together, our results indicate that mitochondrial dysfunction induces IAA production and activates IAA conjugation, which are both greatly enhanced in the *oxt6-1* mutant, suggesting that CPSF30 exerts negative regulatory roles in these processes. In addition to the increase in IAA contents, both ABA and ACC levels were higher, but tZR levels were much lower, while GA24 remained constant in the *oxt6-1* mutant relative to the wild type, again suggesting negative effects imposed by CPSF30 on ABA and ACC accumulation. Interestingly, the basal transcript levels of phytohormone-responsive genes are generally relatively low; as a result, the AA-induced activation of phytohormone responses was strongly enhanced in the *oxt6-1* mutant (Figure 8). The dual effects of AA-induced mitochondrial stress on the auxin pathway can be reflected by 1) the AA-induced activation of IAA biosynthesis and conjugation and 2) the upregulation and downregulation of transcription and translation of auxin-responsive genes, which is attributed in part to AA-induced 3′ UTR shortening. Given that *yuc* and *pin* mutants were more sensitive to AA-induced growth inhibition, we propose that activation of auxin production and the maintenance of auxin homeostasis enhances plant fitness under mitochondrial stress conditions. Cell wall integrity is another critical determinant in plant growth and stress responses via retrograde regulation and/or modulation of plant hormones (Hu et al., 2016; Zhao et al., 2021). Here, we showed that the expression of cell wall–related genes indeed responds to mitochondrial stress co-transcriptionally via APA in addition to transcriptional responses. Notably, most cell wall–related transcripts whose 3′ UTR underwent shortening had high TE, which likely counteracts the downregulation of their transcript levels upon AA treatment (Figure 2E, Figure S2D), suggesting that 3′-end processing acts as a mechanism to reach a cellular balance for cell growth by controlling cell wall integrity and/or expansion under stress conditions. Cell wall integrity is also a critical determinant in plant growth and stress response (Wolf et al., 2012; Hu et al., 2016; Zhao et al., 2021). It is not surprising that mitochondrion-to-nucleus communication resets growth and fitness through cell wall integrity signaling. Recent studies have also revealed that the interplay between the cell wall and phytohormones mediates cell growth and stress tolerance (Aryal et al., 2020; Zhao et al., 2021). Given the insensitivity of cell wall mutants to AA-induced growth inhibition (Figure 11), we propose that defects in cell wall integrity confer mitochondrial stress tolerance.

Here, we present evidence that mitochondrial stress induces global nuclear mRNA 3′ UTR shortening. The histone demethylase JMJ30 controls the choice of PAS in part via co-action with cleavage and polyadenylation specificity factor CPSF30. JMJ30- and CPSF30-regulated APA reprogram nuclear gene expression in response to mitochondrial perturbation, thus bridging mitochondrion-to-nucleus communication to cope with stress for shaping plant growth.

## Materials and methods

### Plant materials and growth conditions

All wildtype, mutant, and transgenic Arabidopsis (*Arabidopsis thaliana*) lines used in this study were in the Columbia (Col-0) accession. The *oxt6-1* (SALK_049389) and *jmj30-1* (SAIL_811_H12) mutants were obtained from the Arabidopsis Biological Resource Center (ABRC). The *yuc9*, *yuc1 yuc2 yuc4 yuc6*, *pin1-11*, and *pin2* mutants are gifts from Dr. Yan Zhao, and *cesa3*, *cesa6*, *mur1*, and *mur4* are gifts from Dr. Chunzhao Zhao. All Arabidopsis seedlings used for the analyses of the molecular and growth phenotypes were grown on half-strength Murashige and Skoog (MS) medium with 1% sucrose or on MS plates with 0.8% agar or 0.6% Phytagel supplemented with or without stressors. Plates were placed in growth chambers at 22°C under 100 μmol m^−2^ s^−1^ light (WT5-LED14, FSL) in a 16-h-light/8-h-dark photoperiod.

### Transgene construction and transformation

The full-length *CPSF30* (At1g30460.3, short version of CPSF30) and *JMJ30* (At3g20810.1) coding sequences were amplified by RT-PCR and cloned in-frame into the binary vectors pEARLEYGATE104 and pEARLEYGATE203 to obtain *35S:YFP-CPSF30 and 35S:cMyc-JMJ30.* A *CPSF30* (At1g30460.3) genomic fragment containing a 1.5-kb of promoter sequence upstream of the ATG was amplified by PCR and cloned in-frame into the binary vector pEARLEYGATE303 to obtain *pCPSF30:gCPSF30-cMyc.* The full-length *JMJ30* coding sequence was cloned downstream of the 35S promoter sequence in the modified pCAMBIA1300 vector to obtain *35S:JMJ30-mCherry* for the examination of co-localization with *35S:YFP-CPSF30.* The full-length *CPSF30* and *JMJ30* coding sequences were also cloned in pSITE-nEYFP and pSITE-cEYFP vectors, and in pGBKT7:BD and pGADT7:AD vectors for BiFC assay and yeast two-hybrid assay, respectively. The sequences of the primers used for the amplification of *CPSF30* and *JMJ30* constructed in different vectors are provided in Supplemental Data Set S9. Transgenic plants were generated by transforming *35S:YFP-CPSF30* and *35S:cMyc-JMJ30* into wildtype Col-0 and *pCPSF30:gCPSF30-cMyc* into the *oxt6-1* mutant using the floral dip method (Clough and Bent, 1998) and selected on medium containing 10 μg/mL Basta (phosphinotricin). For the co-immunoprecipitation (Co-IP) assay, the *35S:YFP-CPSF30* construct was introduced into *35S:cMyc-JMJ30* transgenic plants by crossing to obtain plants harboring both transgenes.

### Growth assessment under stress

Arabidopsis seeds were surface sterilized and sown on solidified half-strength Murashige and Skoog (MS) medium plates. After stratification at 4°C for 2 d in the dark, the plates were placed vertically at 23°C, under a long-day photoperiod. Five-day-old seedlings were transferred to MS plates containing 50 µM antimycin A (AA), 10 µM ABA, 100 nM IAA, or 2 mM H_2_O_2_ and returned to long-day conditions at 22°C for 5–10 d before the phenotypes were recorded by taking photographs and measuring primary root length and fresh weight.

### Confocal imaging of protein subcellular localization and BiFC assay

The subcellular localization of CPSF30 and JMJ30 was examined using confocal microscopy (Zeiss 710) in *N. benthamiana* leaf cells transiently infiltrated or co-infiltrated with the constructs *35S:YFP-CPSF30* and *35S:JMJ30-mCherry.* YFP and chlorophyll were excited at 514 nm, and the resulting fluorescence was detected over the ranges 525–600 and 650–720 nm, respectively. mCherry was excited at 561 nm, and its fluorescence was detected over the range 578–637 nm.

### RNA analyses

Total RNA was extracted from less than 100 mg of 10-d-old seedlings that were grown vertically on half-strength MS plates or treated with 50 μM AA or an equal volume of DMSO for 4 h in liquid MS medium using RNAprep Pure Plant Plus Kit (TIANGEN, DP441). First-strand cDNA was synthesized using a HiScript^®^ II cDNA Synthesis Kit (Vazyme R212-02) and oligo d(T) for nuclear genes. qPCR was conducted with iTaq Universal SYBR^®^ Green Supermix (BIORAD) on a Roche 480 Real-Time PCR system following the manufacturer’s instructions. The expression levels of each gene were normalized to those of *ACT2* as an internal control.

### Poly(A) Tag library construction and sequencing

Wildtype (Col-0), *jmj30-1*, and *35S:cMyc-JMJ30* seedlings were grown vertically on half-strength MS medium plates. Ten-day-old whole seedlings were then transferred into nine-well plates containing half-strength liquid MS medium overnight with shaking in the dark, before being treated with 50 µM AA or an equal volume of DMSO for 4 h. Total RNA from AA-treated and DMSO-treated seedlings was extracted using Trizol reagent (Invitrogen). The poly(A) tag libraries were constructed using the methods described in a previous study (Wang et al., 2017). Briefly, 2 µg of high-quality total RNA (RNA integrity number > 8) was fragmented in the presence of Mg^2+^ in SMARTScribe Reverse Transcriptase buffer (Clontech, Cat. 639537), and mRNA fragments with a poly(A) tail were isolated by Oligo d(T)_25_ Magnetic Beads (NEB Biolab, S1419S). Illumina sequence adaptors were added during reverse transcription with SMRT methods (Wang et al., 2017). The cDNAs were amplified and size-selected to obtain final PAS-seqlibrary. The final libraries were sequenced on an Illumina HiSeq 2500 platform as previously described (Wang et al., 2017), generating 150-bp single-end PAS-seqreads and extracted for downstream bioinformatics analysis. The RNA-seq libraries were constructed with a TruSeq RNA Library Prep Kit v2 (Illumina Inc, San Diego, US, Cat. RS-122-2001) and sequenced on an Illumina NovaSeq 6000 platform to generate paired-end 150-bp reads as previously described (Ye et al., 2020).

### Bioinformatics analysis of poly(A) sites

The bioinformatics workflow of polyadenylation was documented in a previous study (Ye et al., 2021). Briefly, the PAS-seqreads were mapped to the Arabidopsis TAIR10 genome using STAR (v2.7.5c) with default parameters (Dobin et al., 2013). The newly developed R package QuantifyPoly(A) (Ye et al., 2021) with default parameters was used to remove internal priming and cluster poly(A) sites within 24 nt of poly(A) clusters (PACs) (Wu et al., 2011a). Differentially abundant PACs (DAPACs) were identified with the R package DESeq2 (v1.34.0) with the threshold *P*-adj < 0.05 and |Log_2_(fold-change)| > 1 between two conditions (Love et al., 2014). The shift of proximal to poly(A) sites between two conditions was calculated based on the following equation formula: (AA^proximal poly(A) site read1^/CK ^proximal poly(A) site read1^) / (AA^distal poly(A) read2^/CK^distal poly(A) read2^) as described previously (Wang et al., 2017). Variation in 3′ UTR length and 3′ UTR index was calculated with the same strategy described previously (Lin et al., 2020a). The read coverage of single genes was visualized in IGV browser window (Thorvaldsdóttir et al., 2013). The Gene Ontology analysis was performed with the R package clusterprofile (Yu et al., 2012).

### RNA-seq and data analysis

Wildtype (Col-0), *oxt6-1*, *jmj30-1*, and *jmj30-1 oxt6-1* seedlings were grown vertically on half-strength MS medium plates. AA treatment and total RNA extraction were performed as described above. Sequencing libraries were generated from 1 µg total RNA per sample using the NEBNext® Ultra™ RNA Library Prep Kit for Illumina® (NEB, USA) following the manufacturer’s recommendations. Index codes were added to attribute sequences to each sample. The reads were then mapped against the TAIR10 reference genome (https://www.arabidopsis.org). Feature counts was used to count reads for each annotated gene (Liao et al., 2014). R package -DEseq2 were used to normalize and identify differential expression genes with criteria: |log_2_Fold Change| >1 and adjusted p-value < 0.05 (Love et al., 2014). Three biological replicates for each treatment were used for RNA-seq analysis. The PANTHER website (http://pantherdb.org) was used for GO term enrichment analysis of upregulated or downregulated genes (fold-change > 2, Benjamini-Hochberg adjusted *P*-value < 0.01) in RNA-seq data from Col-0, *oxt6-1*, *jmj30-1*, and *jmj30-1 oxt6-1* seedlings treated with 50 µM AA. ImageGP was used to generate GO enrichment plots and heatmaps (Chen et al., 2022).

### Polysome profiling and TE calculations

Sample preparation, collection, and AA treatment were the same as described above for PAS-seqand RNA-seq analyses. Ten-day-old whole seedlings (∼200 mg) were ground in liquid nitrogen and extracted in 1 mL polysome buffer (400 mM Tris-HCl, pH 8.4, 100 mM MgCl_2_, 200 mM KCl, 0.5% [v/v] NP-40, 50 µM EGTA, 50 µg/mL cycloheximide, 50 µg/mL chloramphenicol, and 400 U/mL recombinant RNasin [RNase inhibitor, Promega]). The samples were centrifuged at 12,000g for 10 min at 4°C to remove debris. The supernatant for each sample (1 mL) was loaded on top of a sucrose gradient. Gradients of 20–50% sucrose (11 mL) in gradient buffer (40 mM Tris-HCl, pH 8.4, 10 mM MgCl_2_, and 20 mM KCl) were prepared in polypropylene (14 × 89 mm) centrifuge tubes (Seton, CAN). The gradients were ultra-centrifuged for 3 h at 175,000*g* at 4°C in a SW41Ti rotor (Beckman-Coulter, USA) and analyzed by measuring the absorbance continuously at 260 nm using a piston gradient fractionator (Biocomp Instruments). Total RNA from the polysome or monosome fractions was precipitated and pooled for RNA-seq analysis. Data analysis was performed as described previously (Molenaars et al., 2020).

Translation efficiency (TE) was calculated as the ratio between the transcripts in the polysome fractions (highly translated ribosome-mRNA fractions) and the transcripts in the monosome fractions (lower translated ribosome-mRNA fractions). TEs were then compared between AA-treated samples (AA) and untreated samples (CK). TE was defined as: Log_2_ [(AA^polysome^/CK^polysome^) / (AA^monosome^/CK^monosome^)]. Monosome and polysome fractions were collected as indicated in Figure 2A. Means of three replicates were used to calculate the ratios (See also Supplemental Data Set S2).

### Affinity purification and LC-MS/MS analysis

Ten-day-old seedlings were harvested for isolating total protein. The ground materials were homogenized in 10 mL of lysis buffer (50 mM Tris-HCl, 150 mM NaCl, 5 mM MgCl_2_, 10% [v/v] glycerol, 0.1% [v/v] NP-40, 0.5 mM DTT, 1 mM PMSF, and 1 protease inhibitor cocktail tablet/50 mL, pH 7.5). Following centrifugation, each supernatant was incubated with 50 µL of Anti-cMyc agarose beads (Chromotek, 004282-07-02) at 4°C for 4 h. The resin was washed twice for 5 min with 10 mL of lysis buffer and then five times for 5 min with 1 L of lysis buffer. The proteins bound to the cMyc beads were subjected to on-bead digestion with trypsin. The peptides were sprayed into an Easy-nLC 1200 (Thermo Fisher Scientific) that was coupled online with an Orbitrap Fusion mass spectrometer (Thermo Fisher Scientific). The raw files were searched against the TAIR10 database using Proteome Discoverer.

### Yeast two-hybrid assay

For yeast two-hybrid analysis, the full-length coding sequences of *CPSF30* and *JMJ30* were cloned into the pGBKT7:BD and pGADT7:AD vectors, respectively. The yeast strain Y2H was co-transformed with the corresponding pGADT7 and pGBKT7 constructs and grown on synthetic defined (SD) medium lacking Trp and Leu (SD –Trp –Leu) for selection of transformants. The positive colonies were incubated in SD –Trp –Leu liquid medium at 28°C, and serial dilutions were spotted onto SD – Trp –Leu –His medium plates containing 40 µg/mL X-α-gal.

### Co-IP and gel filtration

Ten-day-old seedlings harboring both the *cMyc-JMJ30* and *YFP-CPSF30* transgenes were used to determine whether JMJ30 interacts with CPSF30 by Co-IP and by gel filtration. For Co-IP, 1 g of seedlings was ground in liquid nitrogen, and the resulting powder was resuspended in 0.5 mL of extraction buffer (50 mM HEPES, 100 mM NaCl, 10 mM EDTA, 10% [v/v] glycerol, 0.2% [v/v] NP-40, 0.5 mM DTT, 1 mM PMSF, and 1 protease inhibitor cocktail tablet/50 mL, pH 7.5) and centrifugation at 12,000g for 20 min at 4°C. Anti-cMyc agarose beads (Chromotek, 004282-07-02) were added to the protein extracts and incubated at 4°C for 4 h. The agarose-bound proteins were precipitated by centrifugation at 12,000g for 1 min at 4°C and washed four times with extraction buffer, boiled in 1× SDS sample buffer, and run on an SDS-PAGE gel for immunoblotting. For gel filtration, 1 g of seedlings was ground in liquid nitrogen and resuspended in 1 mL of lysis buffer (50 mM Tris-HCl, 150 mM NaCl, 5 mM MgCl_2_, 10% [v/v] glycerol, 0.1% [v/v] NP-40, 0.5 mM DTT, 1 mM PMSF, and 1 protease inhibitor cocktail tablet/50 mL, pH 7.5) and centrifuged at 12,000g for 10 min at 4°C. The supernatants were then passed through a 0.22-µm filter, and 500 µL was loaded onto a Superose 6 10/300 GL column (GE Healthcare, 17-5172-01); 500-µL fractions were collected. A 15-µL volume of every fraction was run on a 10% SDS-PAGE for immunoblot assay of cMyc-JMJ30 and YFP-CPSF30. The size of the protein complex was determined with standard proteins. Anti-GFP (TransGen, HT801) and anti-Myc antibodies (TransGen, HT101) were used in immunoblotting for both the Co-IP and gel filtration assays.

### ChIP-PCR assay

The association of H3K27me3 with chromatin was determined using a ChIP-PCR assay as described previously (Saleh et al., 2008; Kaufmann et al., 2010). Ten-day-old seedlings were subjected to crosslinking in 1% (w/v) formaldehyde under vacuum. Thereafter, the nuclei were extracted from the seedlings in nuclei isolation buffer (0.25 M sucrose, 10 mM Tris-HCl, 10 mM MgCl_2_, 1% [v/v] Triton X-100,14.3 M β-mercaptoethanol, 0.1 mM PMSF, and 1 protease inhibitor cocktail tablet/10 mL) and sonicated for chromatin extraction. For H3K27me3 ChIP, an anti-H3K27me3 antibody (Millipore Sigma, 07-449) was incubated with sonicated chromatin overnight, before being purified with protein A agarose (GE Healthcare, 17-1279-01) followed by IP. The precipitated DNA was analyzed by quantitative PCR. The ChIP-qPCR experiments were performed as two biological replicates, and similar results were obtained. Three technical replicates from a representative experiment are shown. *ACT2* (At3g18780) was used as a control. The primers sequences used for the ChIP-qPCR are listed in Supplemental Data Set S9.

### Analysis of phytohormone contents

Sample preparation, collection, and AA treatment were as described above. Ten-day-old whole seedlings were harvested, immediately frozen in liquid nitrogen, ground into powder using a MIXER MILL MM 400 (Retsch, 30 Hz, 1 min), and stored at –80°C until needed. Tissue powder (50 mg) was weighed into a 2-mL plastic microtube and frozen in liquid nitrogen, dissolved in 1 mL methanol/water/formic acid (15:4:1, v/v/v). A 10-μL internal standard solution (100 ng/mL) was added to the extracts as internal standards (IS) for quantification. The mixture was vortexed for 10 min and then centrifuged for 5 min (12,000g at 4°C). The supernatant was transferred to clean plastic microtubes, followed by evaporation to dryness and resuspension in 100 μL 80% (v/v) methanol before being filtered through a 0.22-μm membrane filter for LC-MS/MS analysis using an UPLC-ESI-MS/MS system (UPLC, ExionLC AD, https://sciex.com.cn/; MS, Applied Biosystems 6500 Triple Quadrupole, https://sciex.com.cn/). Phytohormone contents were analyzed using scheduled multiple reaction monitoring (MRM). Data acquisition was performed using Analyst 1.6.3 software (Sciex). Multiquant 3.0.3 software (Sciex) was used to quantify all metabolites. Unsupervised principal component analysis (PCA) was performed using the statistics function prcomp within R (www.r-project.org). Differentially abundant metabolites between groups were determined by variable influence on projection (VIP) ≥ 1 and absolute Log_2_(fold-change) ≥ 1. VIP values were extracted from Orthogonal Projections to Latent Structures Discriminant Analysis (OPLS-DA) results, which also contained score plots and permutation plots, using the R package MetaboAnalystR.

### Data Availability

The RNA-seq, PAS-seq and Ribo-seq data used in this study were deposited in the NCBI SRA database with BioProject ID: PRJNA794925 (RNA-seq and PAT-seq), and with GEO accession: GSE186418, respectively. See Supplemental Data Set S6 for AGI codes of all genes mentioned in this study.

## Acknowledgements

We thank Dr. Yan Zhao for providing auxin mutant seeds and Dr. Chunzhao Zhao for sharing cell wall mutants.

## Author contributions

X.Z., X.G and L.M. conceived the project. H.J., X.Z., J.L. and X.B. generated transgenic plants, performed molecular biology and biochemistry experiments. L. M and W.Z. conducted PAS-seqand RNA-seq analysis. C.M. and Y. Z. and F.X. assisted with experiments. X.Z., L.M., C.-P.S. and X. G. analyzed the data. X.Z wrote the article.

## Supplemental data

**Supplemental Figure S1. Workflow of PAT-seq. A**, Quality of total RNA. Gel image shows the high quality of total RNA used for PAT-seq. **B**, Workflow of sample preparation, library construction, and PAT-sequencing. **C**, AA treatment induces AOX1a expression but has no effects on chloroplast retrograde marker gene expression. **D**, Lincomycin (Lin) and norflurazon (NF) significantly inhibit chloroplast retrograde marker gene expression. **E**, Fraction of genes with different number of poly(A) sites.

**Supplemental Figure S2. TE and TS heatmap of 3′ UTR shortening genes enriched in biological processes.** Enriched terms are indicated to the left of the heatmap, and gene names are indicated to the right. **A**, cell growth (GO:0016049) and response to auxin (GO:0009733); **B**, mitochondrial inner membrane (GO:0005743) and generation of precursor metabolites and energy (GO:0006091); **C**, cytosolic ribosome (GO:0022626); **D**, plant-type cell wall organization or biogenesis (GO:0071669) and cellular response to ABA (GO:0071215). TE: translational efficiency; TS: transcript. HM: Homeodomain-like superfamily protein; HYP: hypothetical protein; Ub-Cytc: Ubiquinol-cytochrome C reductase iron-sulfur subunit; UOXB8: NADH-ubiquinone oxidoreductase B8 subunit; RPS5: RPS5 domain 2-like superfamily protein. See also Supplemental Data Set S2 for details.

**Supplemental Figure S3. *jmj30-1* is a gain-of-function mutant.** Relative *JMJ30* transcript levels were determined by RT-qPCR. Data are means ± SD (*n* = 3). Student’s *t*-test was performed to determine significance. \*\**P* < 0.01. See also Supplemental Data Set S5 for statistical analysis.

**Supplemental Figure S4. AA-induced global alternative polyadenylation in *jmj30-1* mutant and *JMJ30* overexpression plants. A** and **B**, AA-induced global poly(A) site shift in the *jmj30-1* mutant and *JMJ30* overexpression lines. Scatterplot shows the distal-to-proximal shift (red dots) and proximal-to-distal shift (blue dots) in the *jmj30* mutant (**A**) and the *JMJ30* overexpression line (**B**). **C** and **D**, Venn diagrams showing the extent of overlap between genes whose transcripts undergo AA-induced 3′ UTR shortening (3′ UTR-S) and upregulated genes or downregulated genes in the *jmj30-1* mutant (**C**) or *JMJ30* overexpression lines (**D**). **E-G**, GO enrichment analysis. Biological processes enriched among genes whose transcripts undergo AA-induced 3′ UTR shortening in *jmj30-1* mutant (**E**), *JMJ30* overexpression lines (**F**), and wildtype Col-0 (**G**). Common terms enriched in the three genotypes are underlined in red. **H**, Venn diagrams showing the extent of overlap between genes whose transcripts have shortened AA-induced 3′ UTRs (3′ UTR-S) in Col-0, *jmj30-1*, and *JMJ30OX*. See also Supplemental Data Set S3 for details.

**Supplemental Figure S5. Isolation of the *oxt6-1* mutant. A**, Schematic diagram showing the T-DNA insertion in *CPSF30*. The location of the T-DNA in *oxt6-1* (SALK_049389) is indicated by the triangle. LP, RP, are the primers used for genotyping. **B**, DNA gel image showing PCR amplification with the indicated primers using *oxt6-1* or wildtype (Col-0) genomic DNA as template. **C**, Schematic diagram of the *CPSF30S* and *CPSF30L* mature transcripts. Primer pairs used for RT-qPCR of *CPSF30S* and *CPSF30L* are indicated below the diagrams. **D**, Reduction of relative transcript levels of *CPSF30S and CPSF30L* in *oxt6-1* mutants. The relative transcript levels of *CPSF30S* and *CPSF30L* were determined by RT-qPCR. Data are means ± SD (*n* = 3). Student’s *t*-test was performed to determine significance. \**P* < 0.05, \*\**P* < 0.01. See also Supplemental Data Set S5 for statistical analysis.

**Supplemental Figure S6. Growth assessment of the *jmj30-1oxt6-1* double mutant under stress conditions. A**, Growth phenotype. Seedlings of the wildtype Col-0, *jmj30-1* and *oxt6* single mutants, and *jmj30-1 oxt6-1* double mutant were grown on MS plates alone or containing 50 µM AA, 10 µM ABA, 100 nM IAA, or 2 mM H_2_O_2_. Photographs were taken 7 d after transfer to the treatment conditions. **B** and **C**, Mean root length (**B**) and fresh weight (**C**) of the seedlings shown in (**A**). Data are means ± SD (*n* = 3–9). One-way ANOVA was performed to determine significance. See also Supplemental Data Set S5 for statistical analysis.

**Supplemental Figure S7. Locations of the primers for RSI determination.** Primer pairs for the quantification of the abundance of total transcripts or 3′ UTR transcripts are marked with black lines or red lines, respectively. Dark blue stars mark the positions of proximal poly(A) sites.

**Supplemental Figure S8. Locations of the primer pairs for ChIP-PCR. A**, Schematic diagrams of gene models showing the position of primer pairs used for the quantification of H3K27me3 levels over the promoter region (P1), near the ATG codon (P2), in the last coding exon (P3), and over the 3′ UTR (P4 and P5). Light blue stars mark the positions of proximal poly(A) sites. **B**, ChIP-qPCR analysis of H3K27me3 over the *CPSF30* locus. P1 and P2 regions are shared by *CPSF30S* and *CPSF30L*, and P3 and P4 are specific to *CPSF30S* or *CPSF30L*, respectively.

**Supplemental Figure S9. Heatmap of differentially expressed APA factors in AA-treated and control samples.** Heatmap representation of transcript levels for genes encoding APA factors in AA-treated Col-0, *jmj30*, *oxt6*, and *jmj30 oxt6* double mutant seedlings and their controls.

**Supplemental Figure S10. Relative transcript levels of *CPSF30* and *JMJ30*. A**, Fold-change in relative transcript levels for *CPSF30* and *JMJ30* in the indicated mutants vs wildtype Col-0. The left y-axis is for *CPSF30*, and the right y-axis is for *JMJ30*. **B**, Fold-change in relative transcript levels for *CPSF30* and *JMJ30* in AA-treated vs control samples.

**Supplemental Data Set S1. PAS-seqand RNA-seq data of Col-0.**

**Supplemental Data Set S2. RNA-seq data of the monosomal and polysomal RNA for translation efficiency analysis.**

**Supplemental Data Set S3. PAS-seqand RNA-seq data of *jmj30-1* and *JMJ30OX*.**

**Supplemental Data Set S4. RNA-seq data of Col-0, *jmj30-1*, *oxt6-1* and *jmj30-1 oxt6-1*.**

**Supplemental Data Set S5: Statistics Information.**

**Supplemental Data Set S6. Primer sets and antibodies.**

